# Impact of operational conditions on drinking water biofilm dynamics and coliform invasion potential

**DOI:** 10.1101/2023.09.21.558492

**Authors:** Fien Waegenaar, Cristina García-Timermans, Josefien Van Landuyt, Bart De Gusseme, Nico Boon

## Abstract

Biofilms within drinking water distribution systems serve as a habitat for drinking water microorganisms. However, biofilms can negatively impact drinking water quality by causing water discoloration and deterioration and can be a reservoir for unwanted microorganisms. In this study, we investigated whether indicator organisms for drinking water quality, such as coliforms, can settle in mature drinking water biofilms. Therefore, a biofilm monitor consisting of glass rings was used to grow and sample drinking water biofilms. Two mature drinking water biofilms were characterized by flow cytometry, ATP measurements, confocal laser scanning microscopy and 16S rRNA sequencing. Biofilms developed under treated chlorinated surface water supply exhibited lower cell densities in comparison with biofilms resulting from treated groundwater. Overall, the phenotypic as well as the genotypic characteristics were significantly different between both biofilms. In addition, the response of the biofilm microbiome and possible biofilm detachment after minor water quality changes were investigated. Limited changes in pH and free chlorine addition, to simulate operational changes that are relevant for practice, were evaluated. It was shown that both biofilms remained resilient. Finally, mature biofilms were prone to invasion of the coliform, *Serratia fonticola*. After spiking low concentrations (i.e. ± 100 cells/100 mL) of the coliform to the corresponding bulk water samples, the coliforms were able to attach and get established within the mature biofilms. These outcomes are emphasizing the need for continued research on biofilm detachment and its implications for water contamination in distribution networks.

**Importance:** The revelation that even low concentrations of coliforms can infiltrate into mature drinking water biofilms highlights a potential public health concern. Nowadays, the measurement of coliform bacteria is used as an indicator for fecal contamination and to control the effectiveness of disinfection processes and the cleanliness and integrity of distribution systems. In Flanders (Belgium), 533 out of 18840 measurements exceeded the established norm for the coliform indicator parameter in 2021, however, the source of microbial contamination is mostly unknown. Here, we showed that mature biofilms, are susceptible to invasion of *Serratia fonticola*. These findings emphasize the importance of understanding and managing biofilms in drinking water distribution systems, not only for their potential to influence water quality, but also for their role in harboring and potentially disseminating pathogens. Further research into biofilm detachment, long-term responses to operational changes, and pathogen persistence within biofilms is crucial to inform strategies for safeguarding drinking water quality.

## 1 Introduction

Microbial communities are ubiquitously present in drinking water distribution systems (DWDS). Over 98% of these microorganisms form biofilms on pipe materials or are associated with loose deposits (1–3). Biofilms can contribute to drinking water discoloration, the transformation of organic compounds, the decay of free chlorine, microbial regrowth, the formation of disinfection by-products and unwanted odor compounds, etc. (4–7). Additionally, they are recognized as potential sources of opportunistic pathogens (3, 8–12). Previous researchers detected pathogens such as *Mycobacteria* spp. and *Legionella* spp. as well as fecal indicators such as *Escherichia coli*, in biofilm samples from full-scale distribution networks (9, 11). Moreover, *Pseudomonas aeruginosa*, *Mycobacterium avium* and *Legionella pneumophila* can persist in young biofilms after spiking 10^5^-10^6^ cells/mL (12, 13). As a result, biofilms are a potential risk to human health, as biofilm cells can be released to the planktonic water phase under certain conditions. To ensure the quality of drinking water, the measurement of coliform bacteria serves as an indicator for fecal contamination, playing a pivotal role in assessing the effectiveness of disinfection processes and the cleanliness and integrity of distribution systems (14, 15). In Flanders (Belgium), 533 out of 18840 measurements exceeded the established norm for the coliform indicator parameter in 2021, with a maximum value of 201 coliforms per 100 mL (14). Similar maximum concentrations (e.g., 129 coliforms/100 mL or 175 coliforms/100 mL) have been measured before in other full-scale studies performed in the US and Iran (16, 17). However, the source of microbial contaminations is often not retrievable (14, 18).

Characterizing the microbial compositions and phenotypic attributes of biofilms, especially in the context of drinking water, poses challenges due to the practical difficulties in sampling. Previous researchers used laboratory setups to investigate the development and community compositions of biofilms (19–26). For example, biofilm formation rates, measured in terms of ATP activity, and the deposition of iron and manganese have been studied using biofilm monitors consisting of glass rings (27–29). Relatively new sampling techniques consist of a coupon holder to implement in a part of the drinking water distribution systems (DWDS) or in pilot-scale networks (30, 31). Ginige et al. (28) showed that the microbial ATP content on glass rings was 80% less than that on plastic coupons for young biofilms. However, inert glass allows the bulk water characteristics to be the only variable determining biofilm formation and composition (27).

A typical drinking water biofilm comprises a diverse microbial community attached to distribution pipes and immersed in a self-produced matrix of extracellular polymeric substances (EPS), predominantly composed of polysaccharides and proteins (3, 32, 33). The bacterial biofilm community is dominated by *Proteobacteria*, more specifically *Alpha*- and *Gammaproteobacteria*. Notable genera found in biofilms on distribution pipes include *Pseudomonas*, *Sphingomonas* and *Acinetobacter* (1, 6, 31, 34–41). The biofilm environment provides protection against various environmental challenges, including antibiotics, metals, disinfectants like free chlorine, and changes in operational conditions such as shear stress. This protection is attributed to the presence of the EPS matrix and the interconnected processes among the biofilm bacteria (42–44). The formation of biofilms, along with their corresponding phenotypic structure and the existing microbial community, is primarily influenced by the raw water source and treatment processes (e.g., microbial and nutrient composition) (3, 21, 45–48). For instance, previous studies reported a shift in the microbial biofilm community and concentrations on distribution pipes after switching from drinking water with more carbon and a higher conductivity to less turbid water with a lower nutrient content (46, 47). In addition, the composition of drinking water biofilms, and consequently biofilm detachment, is affected by factors such as water residence time, pipe materials, free chlorine concentrations, and temperature (5, 23, 25, 49–55). Former studies mainly focused on the bacterial removal effectiveness of hydrodynamic stressors such as flushing, increasing chlorination concentrations, or interrupted water flow (24, 50, 52, 55). For example, shock chlorination (10 mg Cl_2_/L, 60 minutes contact time) has been shown to remove 75% of the biofilm bacteria (24). However, there is a notable gap in the literature regarding the impact of minor operational variations, commonly encountered in practice, on the dispersal of biofilms into the water phase.

Here, two biofilm monitors, consisting of glass rings were set up to investigate whether unwanted coliforms can intrude into mature drinking water biofilms. First, the biofilms were characterized regarding bacterial cell density and community composition, as well as phenotypic parameters such as biofilm biomass and thickness. Secondly, the response of the biofilm microbiome and possible biofilm detachment after minor water quality changes were investigated. These quality changes (pH, free chlorine concentration) were specifically chosen to simulate operational changes that are relevant for the full-scale DWDS. Finally, the survival of the coliforms in the bulk water phase was evaluated, and attachment of these coliforms on the biofilm was investigated using confocal laser scanning microscopy and qPCR. We chose to spike coliforms in low concentrations (∼100 cells/100 mL), in order to simulate real-life contaminations and thus conditions relevant for practice.

## 2 Materials and Methods

### Biofilm sampling device, conditions and experimental design

A KIWA biofilm monitor was used (KWR, The Netherlands) consisting of 38 glass rings to collect drinking water biofilm (Figure A1) (27). This monitor was placed at two different locations for 17 months, receiving a continuous flow of 270 L/h, according to the manufacturer’s instructions. Monitor 1 was placed at the outlet of a drinking water reservoir receiving treated groundwater, before the UV post-disinfection step (Table A1). Monitor 2 was placed at the outlet of a drinking water tower receiving treated surface water with residual free chlorine. Water quality parameters were measured by the respective drinking water providers (56). The total organic carbon (TOC) concentration was measured at the end of the experiment (18). A timeline of the experiments conducted is represented in Table 1 and briefly described below.

**Table 1:** Experimental design detailing timepoints, specific experiments, water flow, number of rings analyzed, and types of analyses conducted. To perform FCM, ATP, 16S sequencing, and qPCR, a ring was placed in 10 mL of autoclaved tap water and subjected to sonication (3 cycles of 2 minutes each) to detach the biofilm.

| Timepoint (months) | Place | Experiment | Water flow (L/h) | Rings taken for analysis | Type of analysis |
| --- | --- | --- | --- | --- | --- |
| 1 | Water tower/reservoir | Growing biofilm | 270 | / | / |
| 11 | Water tower/reservoir | Growing biofilm | 270 | 4 | FCM, ATP, 16S |
|  |  |  |  | 3 | CLSM (DAPI) |
| 11 | Water tower/reservoir | Growing biofilm | 270 | 6 | CLSM: test EPS staining |
| 17 | Water tower/reservoir | Growing biofilm | 270 | 4 | FCM, ATP, 16S |
|  |  |  |  | 3 | CLSM (DAPI, EPS) |
| 18 | Lab | Changing conditions: LSI = -0.50 | 42 | Hour 0: 2 | FCM |
| 18 | Lab | Changing conditions: LSI = 0.30 | 42 | Hour 0: 2 | FCM |
| 19 | Lab | Changing conditions: HOCl addition | 42 | Hour 0: 2 | FCM |
| 20 | Lab | Invasion (1<sup>st</sup> experiment) | 42 | Day 0: 2 | FCM, qPCR, CLSM (DAPI) |
|  |  |  |  | Day 4: 2 | FCM, qPCR, CLSM (DAPI) |
|  |  |  |  | Day 7: 1 | CLSM (DAPI) |
| 24 | Lab | Invasion (2<sup>nd</sup> experiment) | 42 | Day 0: 2 | FCM, qPCR, CLSM (DAPI) |
|  |  |  |  | Day 8: 1 | qPCR, CLSM (DAPI) |

After 11 months and 17 months, 3 rings were analyzed with confocal laser scanning microscopy (CLSM). 4 rings were put in 10 mL autoclaved tap water and sonicated in a water bath (37 kHz, Elma – Ultrasonic, Belgium) 3 times for 2 minutes with a vortex step in between according to the manufacturer’s instructions, to detach the biofilm. The biomass was quantified in terms of ATP and total cell counts (TCC) and the microbial community was characterized with 16S rRNA amplicon sequencing as described further. In the second part of the study, the KIWA monitors were transported to laboratory conditions to be able to manipulate the water conditions and to perform invasion experiments on the 17 months old biofilms. During these experiments, the corresponding water types were used (Table A1). The monitor was connected to a 10 L plastic vessel and the water was pumped (WM 323 peristaltic pump, Watson Marlow, Belgium) and recirculated over the biofilm monitor. The pump was operating on 150 rpm and the flow was 42 L/h. The experiments were performed at room temperature (i.e., 22 ± 2°C). As the biofilms were consistently exposed to varied experimental conditions, flow cytometry was conducted on a ring before each experiment to define the total cell density (Table A2).

### Changing operational conditions in terms of the Langulier Saturation Index and HOCl addition

The LSI was calculated according to WAC/III/A/011 (57) (Table 1). The pH, conductivity and calcium concentration were measured using a Multi-parameter analyser C1010 (Consort, Belgium), a Hanna Edge Conductivity Meter (HANNA Instruments, Belgium) and the Total Hardness Test (Merck, Belgium), respectively. The alkalinity of the water samples was determined based on WAC/III/A/006 (58). Using 1M NaOH and 1M HCl (Chem-lab, Belgium), the pH was adapted to the a higher and lower Langelier Saturation Index (LSI) (Table 2). In addition, HOCl solution was added to have a free chlorine concentration of 0.20 mg/L and 0.28 mg/L, for the groundwater and surface water samples, respectively. The chlorine concentrate solution was prepared by addition of a NaOCl tablet (B-Care Chemicals, Belgium) to 1 L ultrapure water (Milli-Q, Merck Millipore, Germany). The amount of free chlorine was quantified with the Pocket Colorimeter II (Hach,Belgium).

**Table 2:**
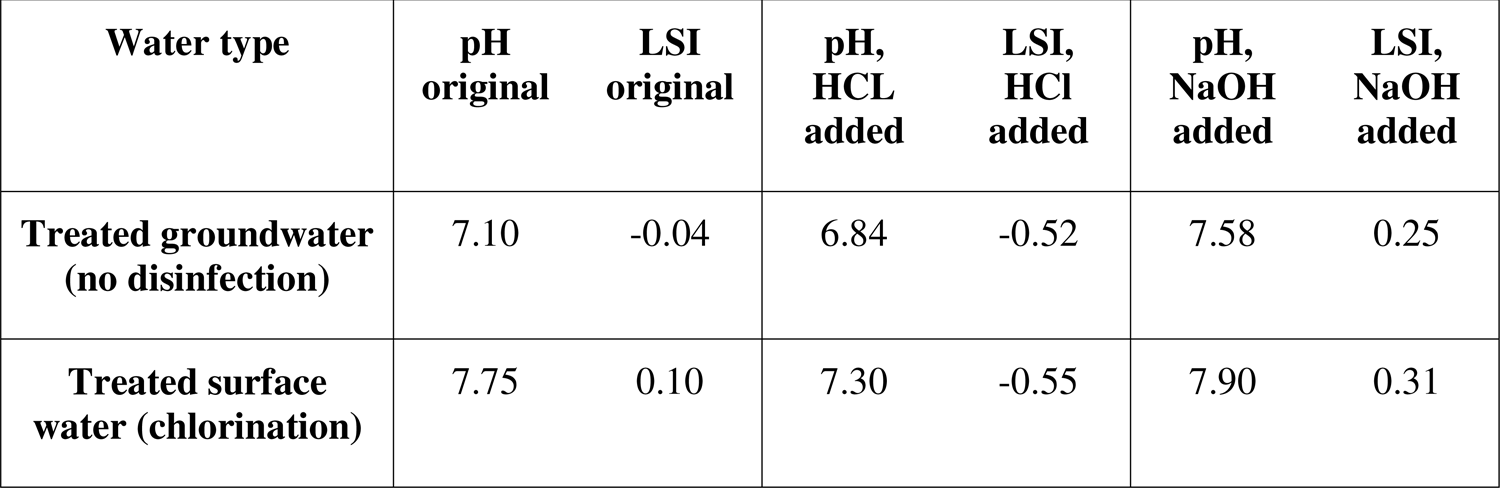
Original pH and corresponding Langelier Saturation Index (LSI) for each water type. To evaluate biofilm detachment, the LSI was changed by decreasing or increasing the pH using HCl and NaOH, respectively.

### Invasion experiments with GFP-labeled *Serratia fonticola*

The strain was isolated from the Flemish drinking water distribution network (Antwerp, Pidpa) and identified with MALDI-TOF mass spectrometry using a Vitek MS (bioMérieux, Marcy-l’Étoile, France) and 16S rRNA gene Sanger sequencing as described by Kerckhof et al. (59) and identified using the NCBI BLAST tool (60). The strain was made ampicillin resistant by serial selections on Luria Broth (LB) agar (Carl Roth, Belgium) with increasing concentrations ampicillin (5 µL/mL to 100 µL/mL) (Merck, Belgium). Each time, the cultures were incubated for 24 hours at 28°C. GFP-labelling was performed using triparental mating. Briefly, the donor strain *Escherichia coli* DH5α pME6012 containing the plasmid ptac-gfp and the helper strain *Escherichia coli* HB101 containing the plasmid pRK2013 needed for conjugation were grown in LB medium at 37°C for 24 hours with 8 µg/mL tetracycline and 50 µg/mL kanamycin, respectively. The acceptor, *Serratia fonticola*, was grown in LB medium with 50 µg/mL ampicillin at 28°C for 24 hours. Next, the cultures were washed twice. They were centrifuged for 5 minutes at 3000 g, the supernatant was removed and 0.22 µm filtered sterile PBS (PBS tablet, Merck, Belgium) was added. Triparental mating was performed by adding10 µL of each culture on LB agar. Plates were incubated at 28°C for 24 hours. GFP-labeled coliforms were selected for resistance to tetracycline (8 µg/mL) and ampicillin (50 µg/mL) and screened for GFP fluorescence using a dark reader (Clare Chemical Research, USA). The final identification of the pure culture was performed with 16S rRNA gene Sanger sequencing as described in Kerckhof et al. (59) and identified using the NCBI BLAST tool (60).

Two invasion experiments were performed by spiking GFP-labeled *Serratia fonticola* to the corresponding water samples (V = 10 L) in plastic vessels which were connected to the KIWA monitor. Therefore, a few colonies were picked from the agar plate and resuspended in 25 mL of 3 g/L sterile R2A broth medium (Oxoid, England) with tetracycline (8 µg/mL) and ampicillin (50 µg/mL) and the tubes were incubated at 28°C and 100 rpm for 24 hours. Subsequently, the culture was washed using sterile 8.5% NaCl as explained before. Afterwards, the culture was transferred to a diluted liquid medium of 50 mg/L R2A broth medium (Oxoid, England) with tetracycline (8 µg/mL) and ampicillin (50 µg/mL) and incubated at 28°C and 100 rpm for 24 hours. The culture was washed before measuring the TCC with flow cytometry. The culture was diluted using sterile 8.5% NaCl and final spike concentrations were ranging from 50 – 500 cells/100 mL. The concentration of *Serratia fonticola* in the bulk water was determined by filtering (3x 100 mL) on S-Pack filters 0.45 µm (Merck, Belgium) using a filtration unit consisting of six filtration funnels and a Microsart e.jet vacuum pump (Sartorius, Germany) and incubation (18 – 23 h, 37°C) on chromogenic coliform agar (CCA) (Carl Roth, Belgium), according to the ISO 9308-1:2014 method for drinking water (56). As a control measure, the bulk water was filtered for selective plating and a ring was extracted for FCM and qPCR analysis, as well as for CLSM analysis, preceding each invasion experiment (Table 1). Water was refreshed after 4 days of recirculation.

### Confocal laser scanning microscopy

Visual characterization of the biofilms was done with a Nikon A1R confocal microscope (Nikon Instruments Inc., USA), which consists of 4 lasers with in total 7 laser lines (405, 457, 476, 488, 514, 561, 639 nm). After 11 months, 3 biofilm rings were taken for CLSM analysis (Table 1). Prior to analysis, samples were fixed in 4% paraformaldehyde (Merck, Belgium) for 24 hours. Fixated biofilm rings were stored at −20°C in a PBS:EtOH 1:1 solution prior to analysis. First, samples were dehydrated in an increasing ethanol series (3 min each in 50, 80 and 96% (v/v) ethanol). Secondly, nucleic acids were labeled with 3 µM DAPI (Excitation/Emission (Ex/Em): 352/464, Merck, Belgium) for 20 minutes in the dark at room temperature. Samples were washed with cold (i.e., ± 6°C), filtered through 0.2 µm, PBS and air-dried. Finally, Fluoroshield™ mounting medium (Merck, Belgium) was added to prevent bleaching. On each ring (n = 3), 1 stack of horizontal plane pictures (10x/0.30 air objective, 512C×C512 pixels equivalent to 1282.58C×C1282.58Cμm) with a z-step of 4.8 μm was taken at 3 locations (randomly selected) of each biofilm ring. After 17 months, an additional set of 3 rings was employed for CLSM analysis, this time incorporating EPS staining. Fixation, storage, and dehydration were conducted the same way as previously described. Then, the protein content was stained for 30 minutes with FilmTracer™ Sypro™ Ruby Biofilm Matrix Stain (Ex/Em: 450/610, Thermofisher Scientific, Belgium) and the carbohydrates with 240 µM Concanavalin A CF640R (Ex/Em: 642/663, Biotium, USA) for 30 minutes. Nucleic acids were labeled with 3 µM DAPI (Ex/Em: 352/464, Merck, Belgium) for 20 minutes. Staining was performed at room temperature (i.e., ± 22°C) and in the dark, after each staining step a washing step with cold (i.e., ± 6°C), filtered through 0.2 µm, PBS. Finally, the rings were air-dried and Fluoroshield™ mounting medium (Merck, Belgium) was added to prevent bleaching. The selection of these fluorophore pairs was based on studies conducted by Fish et al. (61) and Birarda et al. (62). However, as noted by Birarda et al. (62), it is acknowledged that Concanavalin A can bind to glycoproteins and glycolipids. Nonetheless, the abundance of biofilm matrix polysaccharides is presumed to surpass that of these two components, making them the primary contributors to positive carbohydrate staining. On each ring (n = 3), 1 stack of horizontal plane pictures (10x/0.30 air objective, 1024C×C1024 pixels equivalent to 1272.79C×C1272.79Cμm) with a z-step of 4.8 μm was taken at 3 locations (randomly selected) of the ring. For evaluating invasion of the GFP-labeled coliform (Ex/Em: 488/509), each time a ring before and after were analyzed (Table 1). Fixation, storage, dehydration and DAPI staining were conducted the same way as previously described. On each ring (n = 1), 1 stack of horizontal plane pictures (10x/0.30 air objective, 1024C×C1024 pixels equivalent to 1272.79C×C1272.79Cμm or 40x/0.60 air objective, 1024C×C1024 pixels equivalent to 202.42C×C202.42Cμm) with a z-step of 2 μm was taken at 5 locations (randomly selected) of the ring. All samples were analyzed within 14 days. A construction was made and Nunc™ Glass Bottom Dishes (Thermofisher Scientific, Belgium) were used to fit the biofilm ring under the CLSM. Image stacks were processed in ImageJ and the plugin Comstat2 was used to determine biomass, roughness and thickness of the biofilms (64). An automatic threshold (Otsu’s method) was used for all image processing. Particles stained by DAPI are further reported as cells and/or biomass.

### Biomass analysis

ATP and TCC were determined as described in Waegenaar et al. (18). The ATP concentration was measured using the BacTiter-Glo™ Microbial Cell Viability Assay (Promega, Belgium) and luminescence was measured with the Infinite M Plex, multimode microplate reader (Tecan, Switzerland). TCC was measured using an Attune NxT BRXX flow cytometer (ThermoFisher Scientific, USA) and staining was performed with 1 v% of 100 times diluted SYBR Green I solution (10000x concentrate in DMSO, Invitrogen, Belgium). Biofilm samples were 10 times diluted in 0.2 µm filtered Evian and all samples were measured in technical triplicate.

### Molecular analysis of microbial communities

16S rRNA gene amplicon sequencing was performed on biofilm and bulk samples. Biofilm samples (volume = 15 mL) were filtered using Millipore Express PLUS Membranes (Merck, Belgium) and Polycarbonate syringe filter holder (Sartorius, Germany). MF-Millipore Membrane Filters (Merck, Belgium) and a filtration unit consisting of six filtration funnels and a Microsart® e.jet vacuum pump (Sartorius, Germany) were used to filter bulk water samples. DNA extraction was performed using the DNeasy PowerSoilPro kit (Qiagen, Germany), following the manufacturer’s protocol. PCR amplification was performed according to Van Landuyt et al. (65) (see supplementary methods for details). 10 µL genomic DNA extract was send out to LGC genomics GmbH (Berlin, Germany) for library preparation and sequencing on an Illumina Miseq platform with v3 chemistry (Illumina, USA).

### Quantitative polymerase chain reaction (qPCR) to detect *Serratia fonticola*

QPCR assays were performed using a StepOnePlus real-time PCR system (Bio-Rad, Belgium). Specific primers were used to detect the gfp-gen: Forward (5’-AGTGGAGAGGGTGAAGGTGA-3’) and Reverse (5’-ACGGGAAAAGCATTGAACAC-3’). Reactions were performed in a volume of 20 µL consisting of 10 µL of 2x iTAQ universial SYBR Green supermix (Bio-Rad Laboratories, USA), 2.0 µL DNA template, 0.8 µL (10 µM) of each primer, and 6.4 µL nuclease-free water. Amplification conditions were outlined according to the manufacturer’s instructions. Quantification was done a standard curve based on known concentrations of DNA standard dilutions from 10^7^ copies/µL to 10 copies/µL. Reactions were performed in technical triplicates, a negative and positive control was included.

### Data analysis and Statistics

Data analysis was done in R (66) in RStudio version 4.2.1 (67). The Flow Cytometry Standard (.fcs) files were imported using the flowCore package (v2.12.2) (68). The background data was removed by manually drawing a gate on the FL1-H (green) and FL3-H (red) fluorescence channels as described in Props et al. (68). The xlsx package (v4.2.5.2) was used to analyze the data from the confocal microscopy (70). Illumina data was processed using the DADA2 pipeline (v1.28.0) (71). Taxonomy was assigned using the Silva database v138 (72). Further data analysis was performed using statistical packages such as the phyloseq package (v1.44.0) and the vegan package (v2.6-4) (73, 74). Data visualization was done using the ggplot2 (v3.4.3) and ggpubr (v0.6.0) packages (75, 76). The data generated by MALDI-TOF mass spectrometry was analyzed using the MYLA® software. Statistical analysis was done with the dplyr package (v1.1.2) and the vegan package (v2.6-4) (74, 77).

## 3 Results

### Mature biofilm characterization

A KIWA biofilm monitor was used to grow and sample a drinking water biofilm (Figure A1). The monitor was set up at two distinct locations, each receiving water from different sources: treated groundwater without a disinfection step and treated surface water with a chlorination step. The treated surface water was characterized by elevated mineral content, including aluminum, calcium, and nitrate, leading to increased hardness. In contrast, treated groundwater exhibited a higher total organic carbon content and more total colony counts (Table A1).

After 11 and 17 months of biofilm development, we characterized the biofilms through flow cytometry, ATP analysis, confocal laser scanning microscopy (CLSM) and 16S rRNA sequencing (Table 1, Table 3, Figure 1). The cell densities and ATP concentrations of the groundwater biofilms were 10 times higher than those of the surface water biofilms. Similar observations were made using CLSM, where DAPI staining was used to determine the biofilm biomass, roughness and average thickness. The biofilm derived from treated surface water exhibited increased roughness but decreased average thickness both at the 11-month and 17-month intervals. Notably, the treated groundwater biofilms exhibited reduced roughness and average thickness after 17 months compared to the measurements performed after 11 months. In general, significant statistical differences were observed between the two biofilms for each parameter after growing the respective biofilms for 17 months (Table 3). In addition, using 16S rRNA sequencing, a significant difference in the compositions of the two biofilm communities was observed (ANOSIM, p < 0.05, Figure 1). *Chloroflexi* and *Proteobacteria* were identified as the most dominant phyla in the groundwater biofilm samples, constituting 37% and 35% of the community, respectively. *Chloroflexi* were mainly represented by uncultured and unclassified JG30-KF-CM66 (∼30%) and S085 (∼7%) bacteria, whereas only 1.5% of the groundwater bulk bacteria belonged to this phylum. The bulk community was dominated by *Cyanobacteria* (∼26%) and *Alphaproteobacteria* (∼22%), including families like *Hyphomicrobiaceae* (∼3%) and *Hyphomonodaceae* (∼3%) (Figure A2). These families were also detected in the surface water biofilm samples (i.e., ∼3% and ∼11%, respectively) and to a lesser extent in the corresponding bulk samples (i.e., ∼2% and ∼7%, respectively). More than 65% of the bacteria in the biofilms developed under surface water supply were *Alphaproteobacteria*, more specifically *Acetobacteraceae* (∼12%), *Beijerinckiaceae* (∼8.5%) and *Sphingomonadaceae* (∼ 7%). Furthermore, these bacteria were predominant in the surface water bulk samples (∼70%, Figure A2). To conclude, the phenotypic and genotypic characteristics were significantly different between biofilms derived from treated groundwater and treated surface water.

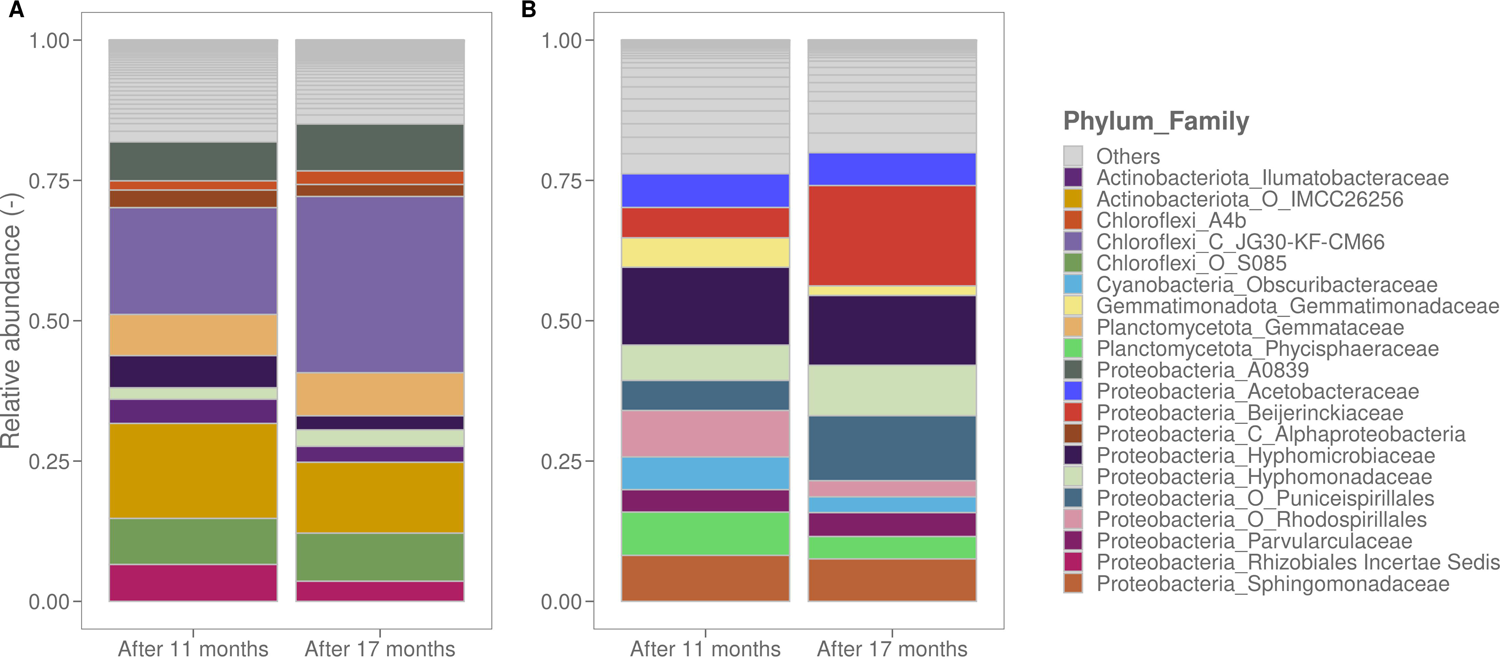

**Table 3:** Characterization of the biofilms using different techniques after 11 and 17 months. Results are presented as an average ± standard deviation. Statistics are done using the Wilcoxon rank sum test or the t test indicated with a ‘t’, and a statistical difference is indicated with a ‘*’. Biofilms resulting from treated groundwater were more dens and active than biofilms resulting treated chlorinated surface water.

| Technique | Parameter | After 11 months |  | P value | After 17 months |  | P value |
| --- | --- | --- | --- | --- | --- | --- | --- |
|  | Water type | Treated groundwater (no disinfection) | Treated surface water (chlorinated) |  | Treated groundwater (no disinfection) | Treated surface water (chlorinated) |  |
| Flow cytometry | Total cell counts (cells/cm<sup>2</sup>) | $(1.01 \pm 0.04) \times 10^7$ | $(5.96 \pm 1.01) \times 10^5$ | $4.11 \times 10^{-5} *$ | $(7.46 \pm 0.58) \times 10^6$ | $(5.23 \pm 3.68) \times 10^5$ | $4.11 \times 10^{-5} *$ |
| ATP analysis | ATP content (pg/cm<sup>2</sup>) | $411.34 \pm 52.58$ | $51.86 \pm 3.22$ | $8.23 \times 10^{-5} *$ | $896.54 \pm 72.11$ | $77.26 \pm 41.48$ | $4.04 \times 10^{-4} *$ |
| CLSM | Roughness (Ra) | $1.25 \pm 0.39$ | $1.97 \pm 0.03$ | 0.06* | $0.78 \pm 0.14$ | $1.68 \pm 0.04$ | $3.97 \times 10^{-5} *$ |
| CLSM | Average thickness ($\mu\text{m}$) | $15.78 \pm 5.89$ | $0.94 \pm 0.57$ | 0.06* | $4.95 \pm 2.62$ | $1.31 \pm 0.33$ | $7.18 \times 10^{-3} *$ |
| CLSM | Biomass ($\mu\text{m}^3/\mu\text{m}^2$) | $5.65 \pm 2.23$ | $0.57 \pm 0.45$ | $0.08^{\text{t}} *$ | $4.10 \pm 3.07$ | $0.25 \pm 0.10$ | $1.59 \times 10^{-4} *$ |
| CLSM EPS | Protein biovolume ($\mu\text{m}^3/\mu\text{m}^2$) | Not determined | Not determined | / | $1.23 \pm 1.09$ | $0.13 \pm 0.07$ | $3.97 \times 10^{-5} *$ |
| CLSM EPS | Sugar biovolume ($\mu\text{m}^3/\mu\text{m}^2$) | Not determined | Not determined | / | $2.73 \pm 2.54$ | $0.29 \pm 0.25$ | $7.94 \times 10^{-5} *$ |

As extracellular polymeric substances (EPS) play a crucial role in the biofilm structure, the protein and sugar content were determined for both biofilm samples after 17 months (Table 1, Table 3, Figure 2). Consistent with the other measurements, the groundwater biofilm exhibited 10 times more EPS biovolume than the surface water biofilm. However, the ratios of sugars to biomass and proteins to biomass were similar for both mature biofilms (Figure A3). The higher bacterial and EPS content in the groundwater biofilm could be attributed to the higher carbon content in the raw water and the absence of disinfectants (Table A1). To illustrate that the biofilms used in further experiments were mature, statistical analyses were performed between the two timepoints, more precisely after 11 and 17 months. Biomass and community composition were selected as key parameters, and no significant differences were observed over time for both biofilms (Table A3).

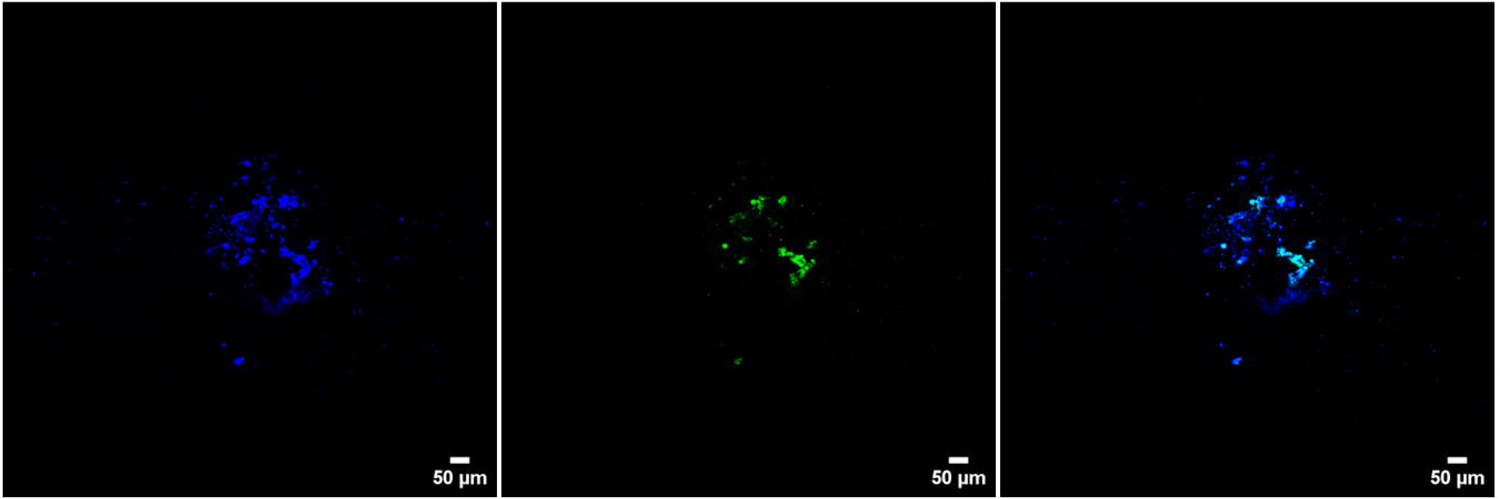

### Effect of operational water quality changes on biofilm detachment

After 17 months of growing a drinking water biofilm, the KIWA monitors were transferred from the water reservoirs to laboratory conditions, and the original water quality was adjusted to examine the response of the biofilm microbiome to minor changes observed in practice (Table 1). More concretely, the effects of slight pH variations and the addition of small concentrations of HOCl on biofilm detachment were investigated. These minor pH changes led to an increase or decrease in the Langelier Saturation Index (LSI) (Table 2). This qualitative index predicts the scale forming potential of water and is based on the measurement of pH, conductivity, alkalinity, calcium ions and temperature (78, 79). Cell counts in the bulk were measured with flow cytometry for 8 hours. Generally, little or no detachment of the biofilm was observed compared to the untreated controls (Figure 3). Linear regression with a confidence interval of 95% was performed to quantify cell release rates to the bulk. If the cell release rate is higher than zero, there is an indication that biofilm cells are dispersed into the bulk water phase. Briefly, detachment was always observed except for two blank conditions. Furthermore, the dispersal of biofilm cells to the bulk water was faster (i.e., higher release rates) when the LSI was changed or when HOCl was added. Therefore, statistics on the residuals and the slopes of the linear regression models was conducted and no significant difference was observed between the controls and the applied operational changes (Table A4).

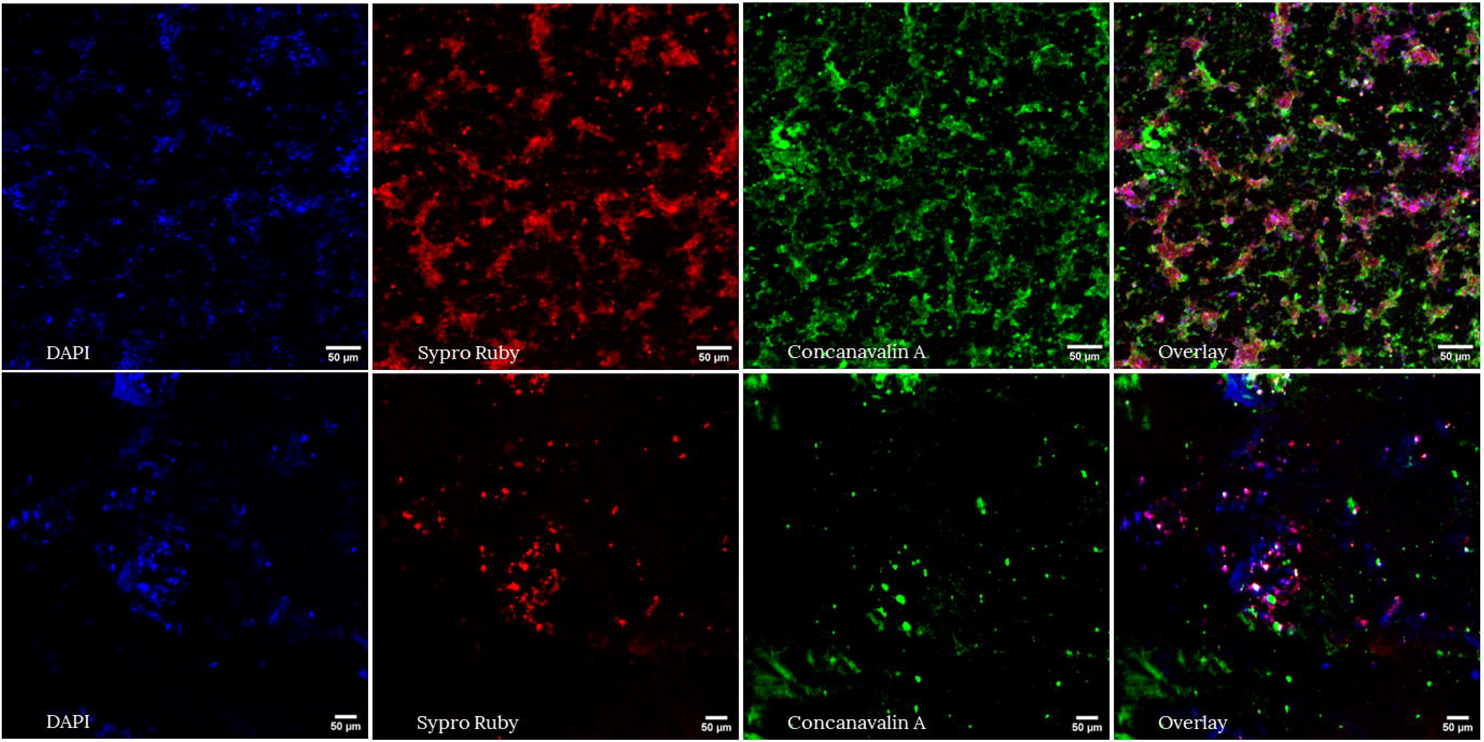

### Invasion of coliforms onto drinking water biofilms

*Serratia fonticola*, isolated from the Flemish distribution network (Belgium), was labeled with a green fluorescent protein (GFP) to distinguish this specific strain from other drinking water biofilm bacteria. It was then introduced into the water supplied to the biofilm monitors. Two invasion experiments were performed for each water type (Table 1). The initial concentration and survival of the coliform in the bulk water were monitored using selective media. QPCR and CLSM were used to determine whether *Serratia fonticola* attached to the biofilm (Table 1). Prior to conducting the invasion experiments, control measures were executed for bulk and biofilm samples, and no coliforms were detected. *Serratia fonticola* was added to the water samples and concentrations were ranging between 15 cells/100 mL and 650 cells/100 mL (Table 4). For treated surface water, the coliforms were still present in the bulk water phase after three hours. After 24 hours, the coliforms were below detection limit (< 0 cells/100 mL) for both water types. Biofilms on the glass rings were analyzed after 4 or 7 days using CLSM and qPCR. For the first invasion experiment, the gene copies were below the limit of quantification (LOQ = 28.57 gene copies/cm²), whereas for the second invasion experiment, 578.59 ± 38.45 and 358.36 ± 24.03 gene copies/cm² of the invader were measured in the groundwater and surface water biofilms, respectively (Table 4). Detection of the GFP-labeled *Serratia fonticola* in the biofilm was also done with CLSM and coliforms were detected 7 or 8 days after the spike for each water type (Figure 4, Figures A4 - A11). Even though mature biofilms seem to be strong microbial ecosystems and minor water quality changes do not lead to major detachment into the bulk phase, they are susceptible to unwanted invasion by coliforms.

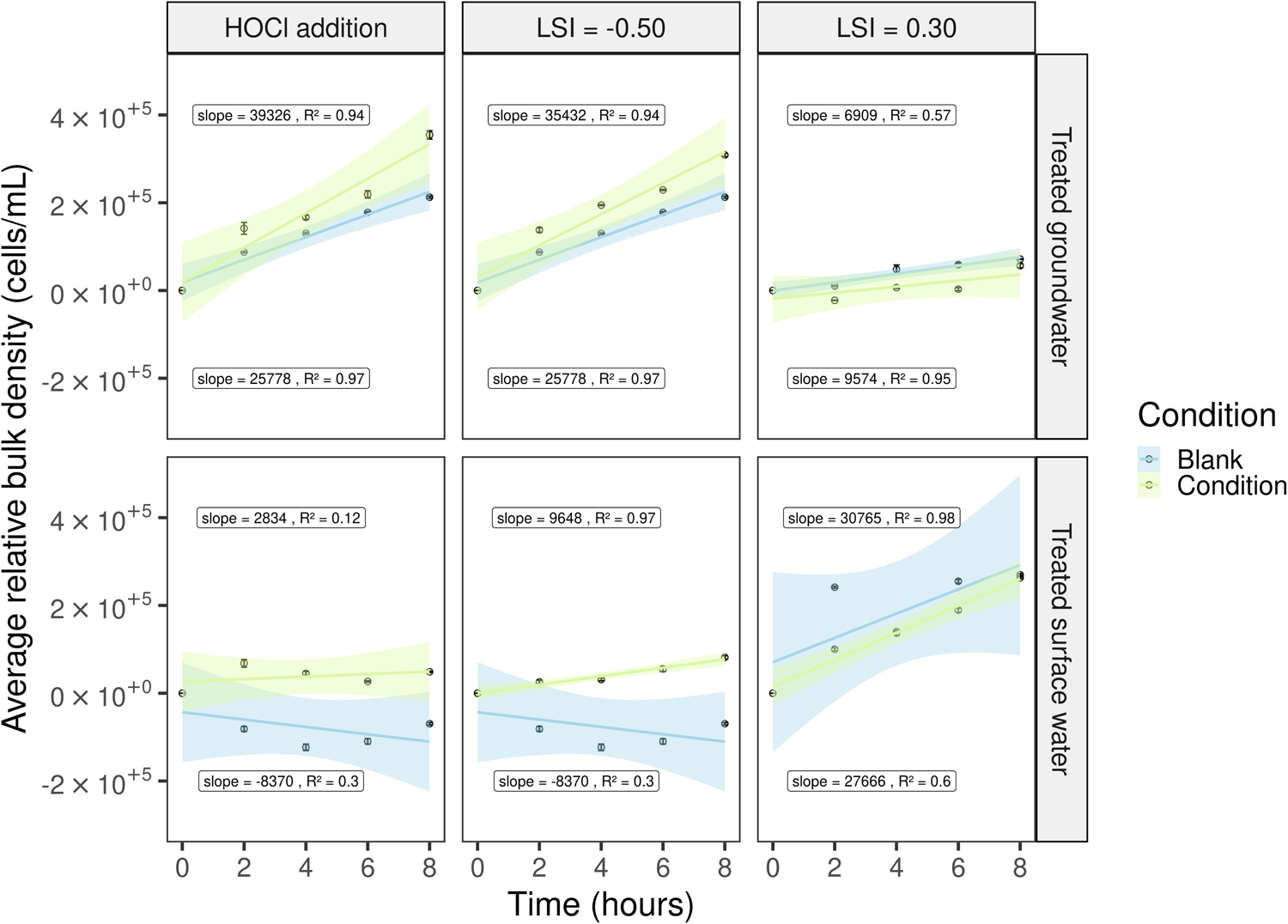

**Table 4:** Two invasion experiments were performed with Serratia fonticola on the mature biofilms. The concentration in the bulk water phase was followed with selective plating (chromogenic coliform agar). Invasion of the coliform in the biofilm was determined using CLSM and qPCR. Positive detection on the respective day after the spike (Dx) with CLSM is indicated with ‘present’. It was not possible to perform CLSM for the groundwater biofilms of the second spike experiment because of calcium precipitation on the rings. Using qPCR, the GFP-labeled coliform was found in the biofilm after 7 days (D7).

| Technique | Experiment | Bulk |  |  | Biofilm |  |
| --- | --- | --- | --- | --- | --- | --- |
|  |  | Chromogenic coliform agar |  |  | CLSM | qPCR |
| Water type |  | Start concentration (cells/100 mL) | After 3 hours (cells/100 mL) | After 24 hours (cells/100 mL) | / | (Gene copies/cm<sup>2</sup>) |
| **Treated groundwater (no disinfection)** | 1<sup>st</sup> invasion experiment | 104.0 ± 10.6 | 0 | 0 | D4: not present<br>D8: present | D4: under LOQ |
| **Treated groundwater (no disinfection)** | 2<sup>nd</sup> invasion experiment | 15.3 ± 3.0 | 0 | 0 | D7: not possible | D7: 578.59 ± 38.45 |
| **Treated surface water (chlorinated)** | 1<sup>st</sup> invasion experiment | 650.0 ± 14.1 | 420.0 ± 17.0 | 0 | D4: not present<br>D8: present | D4: under LOQ |
| **Treated surface water (chlorinated)** | 2<sup>nd</sup> invasion experiment | 39.5 ± 5.0 | 1.0 ± 1.4 | 0 | D7: present | D7: 358.36 ± 24.03 |

## 4. Discussion

### Treated groundwater and chlorinated surface water resulted in significantly different biofilms in terms of community compositions and biomass content

Two biofilm monitors, consisting of glass rings, were employed to investigate drinking water biofilms. They were placed in two distinct reservoirs for 17 months, one receiving treated groundwater without disinfection and the other receiving treated surface water with residual free chlorine. The use of glass as a carrier material ensures reproducible biofilm sampling, considering only the biofilm formation potential. However, due to the absence of nutrient leaching, glass results in lower biofilm density compared to plastic (28). In general, bacterial cell densities of mature biofilms vary from 10^4^ – 10^8^ cells/cm² (2, 3, 22, 80). We observed similar concentrations, although a lower cell density was observed for biofilms developed under surface water supply, probably due to a free chlorine disinfectant (Table 3). Furthermore, the results aligned with previous studies, showing ATP concentrations of 100–1100 pg/cm² for unchlorinated and 10–100 pg ATP/cm² for chlorinated water (38, 81). Regarding CLSM analysis results, lower biomass content was observed for treated surface water biofilms, hovering around 1 µm³/µm², as previously reported in the literature (5). A study by Shen et al. (82) demonstrated a positive correlation between biofilm roughness and adhesion and a negative correlation with detachment. Although significantly higher biofilm roughness was measured for the surface water biofilm, this did not result in higher biomass content. Overall, biofilms from treated groundwater exhibited a 10 times higher FCM cell concentration, biofilm thickness, ATP and biomass content, possibly due to higher cell and TOC concentrations in bulk water and the absence of disinfection (Table 3, Table A1). Furthermore, chlorination not only reduced biofilm density but also decreased EPS production (Figure 2, Table 3) (22). 16S rRNA sequencing revealed a significant difference between treated groundwater and surface water biofilms (Figure 1). The groundwater biofilm community was dominated by *Alphaproteobacteria*, *Chloroflexi* and *Actinobacteriota*, while the surface water biofilm mainly consisted of *Alphaproteobacteria* and *Gemmatimonadota*. Within the *Chloroflexi* phylum, the class JG30-KF-CM66 was most abundant, even though only small concentrations were found in the bulk. This finding is consistent with raw groundwater measurements reported in the literature (Figure 1, Figure A2) (83). Additionally, members of the *Chloroflexi* and *Actinobacteriota* clusters are known to degrade complex organic matter structures that could be present in biofilms (84, 85). On the other hand, previous studies have confirmed that the biofilm and bulk core microbiome of treated chlorinated surface water mainly consists of *Alphaproteobacteria*, such as *Hyphomicrobiaceae* and *Sphingomonadaceae* (20, 26, 53, 86–88). These taxa are both known to easily colonize surfaces, produce EPS, and can degrade a wide range of organic carbon compounds (20, 87, 89).

This study demonstrated that the biofilms entered a mature phase, also referred to as quasi-steady state phase, after 17 months, due to a consistent biofilm density and community during the 11-month to 17-month intervals (Table A3). Previous studies have indicated that this mature phase is characterized by stable cell numbers, EPS formation and maintaining an equilibrium between growth, attachment and detachment (40, 80). Furthermore, we showed that treated groundwater and chlorinated surface water resulted in significantly different biofilms, impacting cell density and community composition. This underscores the importance of both water source and treatment processes in biofilm formation. However, conflicting findings persist regarding the impact of source and treatment in distribution pipes. Previous studies have suggested that the source water mainly shapes the biofilm community composition (36, 41). In contrast, other researchers have shown that there is no significant difference in the biofilm community concerning the drinking water source, and that treatment (e.g., disinfection) is more important for the biofilm core community in distribution pipes (90).

### Mature biofilms react minimally towards operational changes in water quality

While the water quality in drinking water distribution systems generally remains constant, minor operational adjustments can occur, impacting water microbiology (92). For example, additional chlorination because of water works, variations in the quality of raw water sources, mixing of different water types in the DWDS, etc. (91, 92). Previous studies have primarily focused on the influence of severe operational changes, such as flushing and shock chlorination, the goal of this work is to investigate the effects of these minor variations, which frequently occur in practice, on biofilm detachment.

Drinking water providers maintain fixed pH values and free chlorine dosage during treatment to control biofilm formation and corrosion of pipe materials, valves, pumps, etc. (15). To evaluate water corrosivity, the LSI, a qualitative index that predicts the scale forming potential of water, is determined. Both European and Belgian drinking water directives recommend measuring an LSI above −0.5 (14, 93). A slightly negative LSI means more corrosive water that contains carbon dioxide deposits, while a positive LSI indicates CaCO_3_ supersaturated water with the potential to form scale (78). In this study, the LSI of the tested waters was adjusted by changing the pH (Table 2). Additionally, an HOCl solution was added to have a free chlorine concentration of 0.20 mg/L and 0.28 mg/L, for the groundwater and surface water samples, respectively. In general, there was detachment towards the bulk water phase as indicated by the slopes of the regression models being higher than zero. However, no significant biofilm detachment was observed between the untreated controls and the adjusted waters (Figure 3, Table A4). Besides, our drinking water setup, and distribution systems in general, demonstrated resilience (36). The pH was restored after 4 hours, and the free chlorine concentration was below the detection limit (< 0.05 mg Cl_2_/L) after 2 hours of recirculation (data not shown). Similarly, the research of Trihn et al. (5) observed the role of biofilms in free chlorine decay.

However, the implementation of the results should be handled with care, as only the short effect (i.e., 8 hours) of the operational changes was investigated. Continuous pH changes could alter the electrostatic interactions between materials and microorganisms and between microorganisms (8, 83). Former studies have examined the long-term effect of chlorine on biofilms, mentioning that small increases in chlorine concentrations could lead to a decrease in culturable biofilm bacteria and EPS production (42, 52). Furthermore, in our study, biofilms were grown on glass rings, whereas in a full-scale distribution network aged biofilms on iron or plastic piping materials are used, which are more susceptible towards corrosive water. For example, biofilms attached to stainless steel compound pipes are more sensitive to flushing than those attached to ductile cast iron pipes (55).

### Mature biofilms are susceptible towards the invasion of *Serratia fonticola*

In the next part, we added a GFP-labeled coliform, *Serratia fonticola* (i.e., ± 100 cells/100 mL), into the treated groundwater and surface water samples to investigate invasion onto the corresponding biofilms. Water was recirculated over the biofilm monitor for 4 days, and *Serratia fonticola* was followed using selective media (Table 4). Our results indicated that the coliform was unable to survive longer than 24 hours in the bulk waters, possibly because of the oligotrophic drinking water environment or competition with the resident drinking water community (94–97). However, after 7 days, the coliform was detected in the biofilm samples (Figure 4, Table 4). Since confocal microscopy has limitations (e.g., operator dependent) and bleaching of the fluorescent protein was observed, the results were validated using qPCR based on the detection of the fluorescent gene, which was incorporated into the genome of the coliform. In the first invasion experiment, the gene copies were below the limit of quantification, but in the second invasion experiment, detection with qPCR was achieved (Table 4). This variation could be attributed to the sampling day (after 7 days instead of 4 days) or more favorable invasion circumstances. Overall, adhesion and invasion onto a drinking water biofilm depend on nutrient availability, surrounding microorganisms, local hydrodynamics and biofilm roughness (82, 95).

Previous researchers already showed that biofilms could be a reservoir for fecal indicators and pathogens (8, 10). For example, Kilb et al. (98) detected coliforms on rubber-coated valves from distribution networks. Pathogens such as *Pseudomonas aeruginosa* were able to persist in drinking water biofilms after spiking through bulk water samples (12). However, in comparison with the microbial drinking water legislation (i.e., absence in 250 mL and 100 mL for *P. aeruginosa* and total coliforms, respectively) and the observed concentrations in practice (i.e., 200 cells/100 mL), high spike concentrations were used (i.e., 10^5^-10^7^ cells/mL) in former studies (12, 13, 15–17, 95). Here, in this study, we demonstrated that even low concentrations of coliforms (± 100 cells/100 mL) can attach and get established in mature drinking water biofilms. We explicitly chose to use low concentrations as they are relevant for the full-scale practice, where the bigger volumes and flow rates in the DWDS might get contaminated by for example groundwater intrusion after pipe burst, or rain-, river- or wastewater contamination due to wrong connections (99, 100). Furthermore, it is important to notice that *Serratia fonticola* was able to settle in two significant different biofilms regarding cell density and community composition (Figure 1, Table 3, Table 4). As mentioned before, both biofilms were characterized as mature or quasi-stationary. The concept of quasi-stationary phase is introduced by Boe-Hansen et al. (80), who argue that a true stationary phase is never reached in biofilms due to continuous selection influenced by small changes in environmental conditions. This suggests that certain microcolonies within biofilms may be more susceptible at specific moments.

Water stagnation, changes in flow rates, and flushing after water works can disrupt biofilms, potentially releasing coliforms and other unwanted microorganisms into the water phase, leading to contamination and associated health concerns (50, 55, 92). Moreover, detached coliforms may settle elsewhere in the DWDS. Despite these risks, further research about biofilm detachment and effective mitigation strategies for preventing the establishment of unwanted microorganisms is necessary to understand the occurrence of contaminations in distribution networks. Favere et al. (94) propose that pathogens and indicator organisms are considered to be r-strategist (high growth rate at high nutrient concentrations), whereas the naturally drinking water community consist out of K-strategist (high substrate affinity). When producing biostable water through nutrient limitation, the bacterial community is directed towards K-strategists, consequently restricting the survival of the r-strategist. In addition, previous research have indicated that by depleting nutrients (such as carbon, nitrogen or oxygen), both the biological activity in the water and biofilm as well as the production of EPS and subsequent biofilm adhesion, can be reduced (44, 101).

## 4 Conclusion

In conclusion, we have used a biofilm monitor consisting of glass rings to study drinking water biofilms. More specifically, we investigated the response of the biofilm microbiome towards limited operational variations in pH and free chlorine concentration, and the ability of the coliform, *Serratia fonticola*, to settle onto these biofilms. Two mature drinking water biofilms were characterized using several techniques and it was shown that they were significantly different from each other regarding cell density and community composition. To summarize, biofilms resulting from treated groundwater had a 10 times higher cell density, biofilm thickness, ATP and biomass content than biofilms resulting from treated chlorinated surface water. Next, it was observed that the biofilms remained resilient when applying limited changes that are seen in the full-scale drinking water network. Finally, *Serratia fonticola*, spiked at low concentrations through the bulk water phase, demonstrated the ability to attach and get established within the mature biofilms, highlighting the potential for biofilms to act as reservoirs for unwanted microorganisms in drinking water distribution systems.

## Supporting information

Supplemental figures and tables (16S rRNA sequencing results, statistics).

## Acknowledgements

We would like to thank all colleagues of Pidpa, De Watergroep and Farys who were involved in this project. Special acknowledgements go to thank Martine Cuypers (Pidpa), Katrien De Maeyer (Pidpa), Bart Van Calenberge (De Watergroep), Tom Vandermarliere (Farys), Benjamin Buysschaert (EKOPAK), Geert Van Nimmen (Farys) for installing and maintaining the biofilm monitors. Paul Bielen (Pidpa) and Tom Vandermarliere (Farys) for collecting the raw water quality measurements. We thank Katrien De Maeyer (Pidpa) for providing the coliform strain, Jan Roelof van der Meer (Until, Switzerland) for providing the donor and helper strains to label the coliform with a GFP and Jorien Favere (ORB, United Kingdom) for her help in performing the triparental mating experiments. We would like to thank Geert Meesen for his help on the confocal microscope and Tim Lacoere and Inez Roegiers for helping to perform and interpret the qPCR analysis and results. Finally, we would like to thank Jorien Favere and Josefien Van Landuyt to discuss the results of the experiments.

This work was funded by the Research Foundation—Flanders (FWO) (grant number 1S02022N) and by the FWO-SBO Biostable project (grant number S006221N). The work is part of the Ghent University-Aquaflanders Chair for Sustainable Drinking Water, which is supported by Aquaflanders, the federation of Flemish companies that are responsible for drinking water and sewer management (www.aquaflanders.be). F.W., C.G., J.V.L, B.D.G. and N.B. conceived the study. F.W. carried out the laboratory work, analyzed the data, interpreted the results and wrote the paper. C.G., J.V.L, B.D.G. and N.B. interpreted the results and supervised the findings of this work. All authors reviewed and approved the manuscript. The authors declare that there are no conflicts of interest.

## Data availability statement

The datasets presented in this study can be found in online repositories. The names of the repository/repositories and accession number(s) can be found below: https://github.com/waegenaarfien/2023_Unwanted-coliforms-can-hide-in-mature-drinking-water-biofilms.git, https://www.ncbi.nlm.nih.gov/bioproject/PRJNA1015597.

## Supplemental material

Supplemental material for this article may be found at …

