## Supplemental figures and tables (16S rRNA sequencing results, statistics). for "Impact of operational conditions on drinking water biofilm dynamics and coliform invasion potential"

**Running title:**

Coliform invasion in drinking water biofilms.

**SUPPLEMENTARY METHODS**

**16S rRNA amplicon sequencing**

For quality control of subsequent DNA extraction, the PCR amplification of the 16S rRNA gene V3-V4 hypervariable regions was carried out using Taq DNA Polymerase and the Fermentas PCR Master Mix Kit in accordance with the manufacturers’ specifications (Thermo Fisher Scientific, Waltham, MA, USA). The primers used were 341F (5’-CCT ACG GGN GGC WGC AG -3’) and 785Rmod (5’-GAC TAC HVG GGT ATC TAA KCC-3’), with the reverse primer adapted from Klindworth et al. (2013) to enhance coverage. The selection of the V3-V4 hypervariable region of the 16S rRNA gene was based on its optimal characteristics for effective sequencing during 2 x 250 base pair Illumina Miseq sequencing, as suggested by Klindworth et al. (2013). The resulting PCR product, along with the DNA extract and a GeneRuler DNA Ladder Mix (Thermo Fisher Scientific, Waltham, MA, USA), was subjected to electrophoresis on a 2% agarose gel for 30 minutes at 100 V as a control.

**SUPPLEMENTARY FIGURES AND TABLES**

 
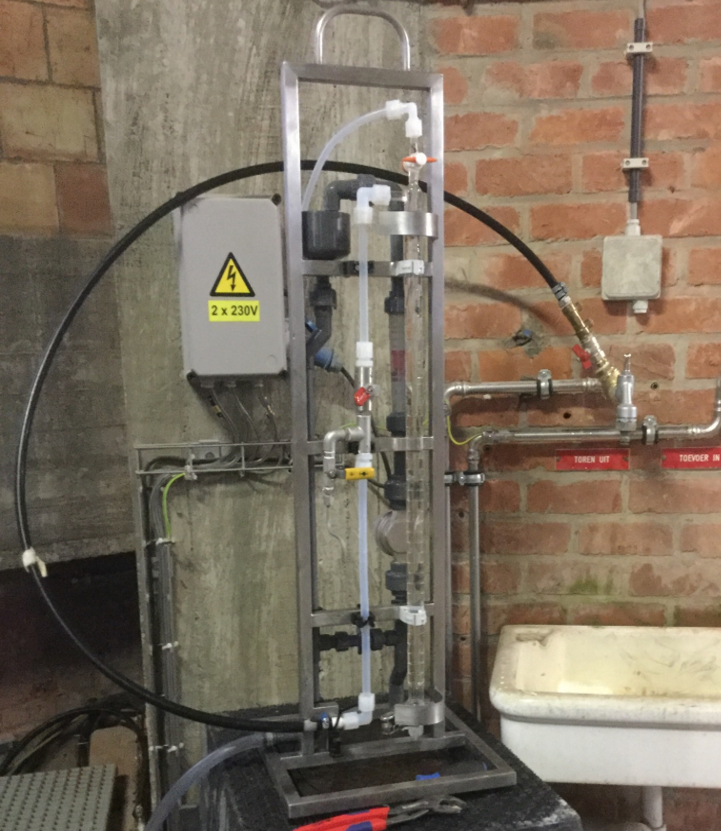

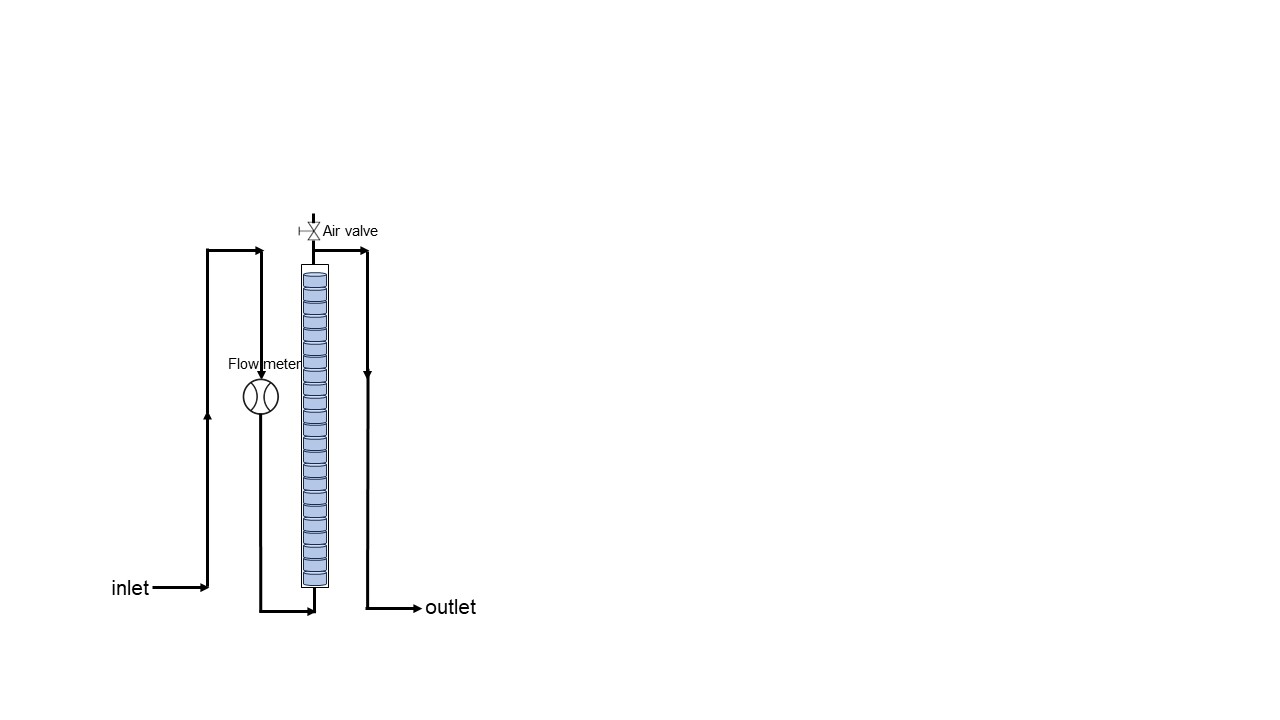

**A**

**B**

Figure A1: **(A)** KIWA monitor consisting of 38 glass rings to collect drinking water biofilm samples. One monitor was placed at a water reservoir receiving treated groundwater whereas the other monitor, located in a water tower, received treated chlorinated surface water. **(B)** Schematic representation of the biofilm monitor, the water flow is indicated with a black arrow and the section with biofilm rings is represented by blue cylinders.

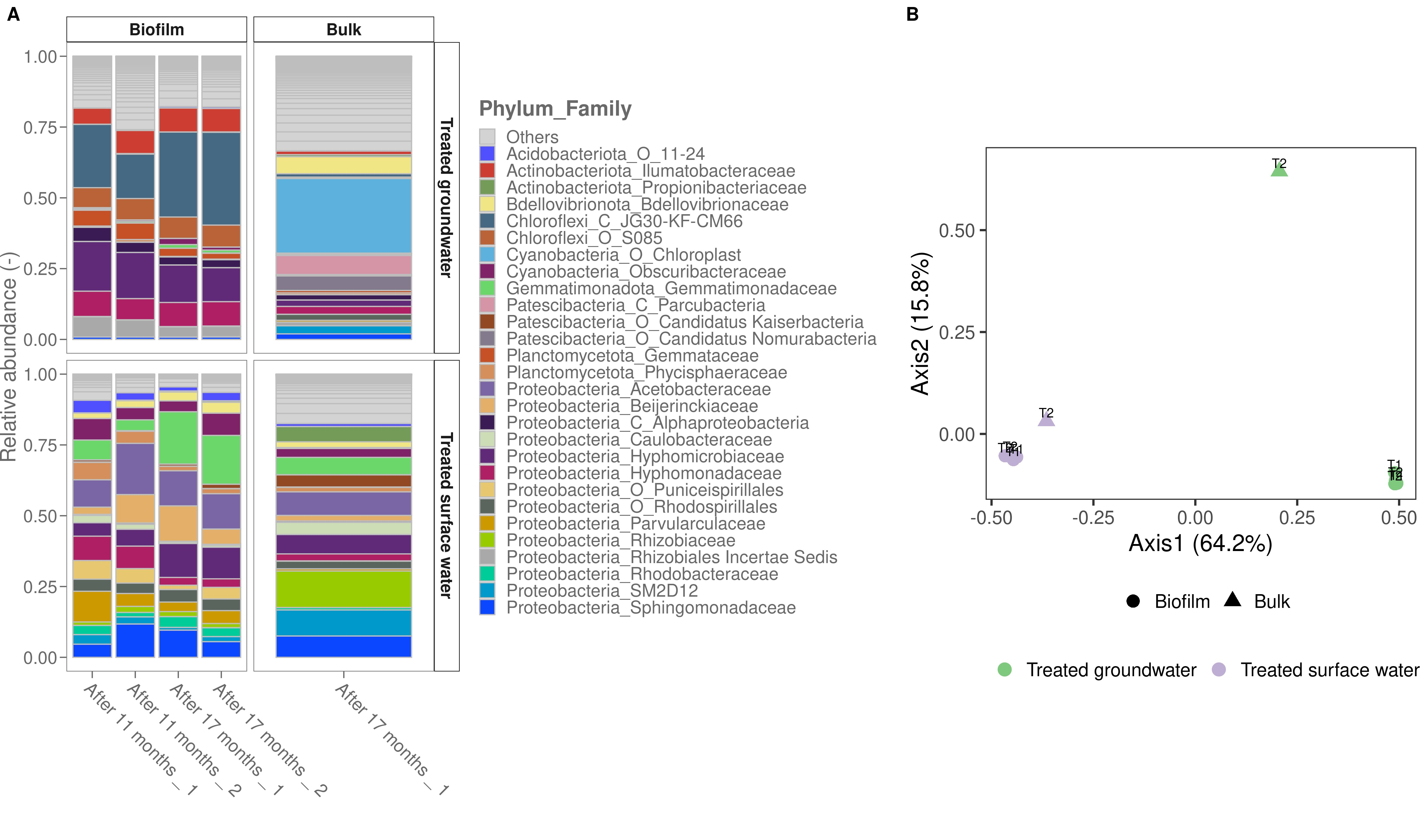

Figure A2: **(A)** Relative abundances of the 29 most abundant families in the biofilm and bulk samples for each water type. Biofilm samples for 16S rRNA sequencing (n = 2) were taken after 11 and 17 months, bulk samples (V = 2L) were taken after 17 months. **(B)** Corresponding PCoA analysis based on the Bray-Curtis distance metric of the sequencing results. The community of bulk and biofilm samples is more similar when considering the surface water samples.

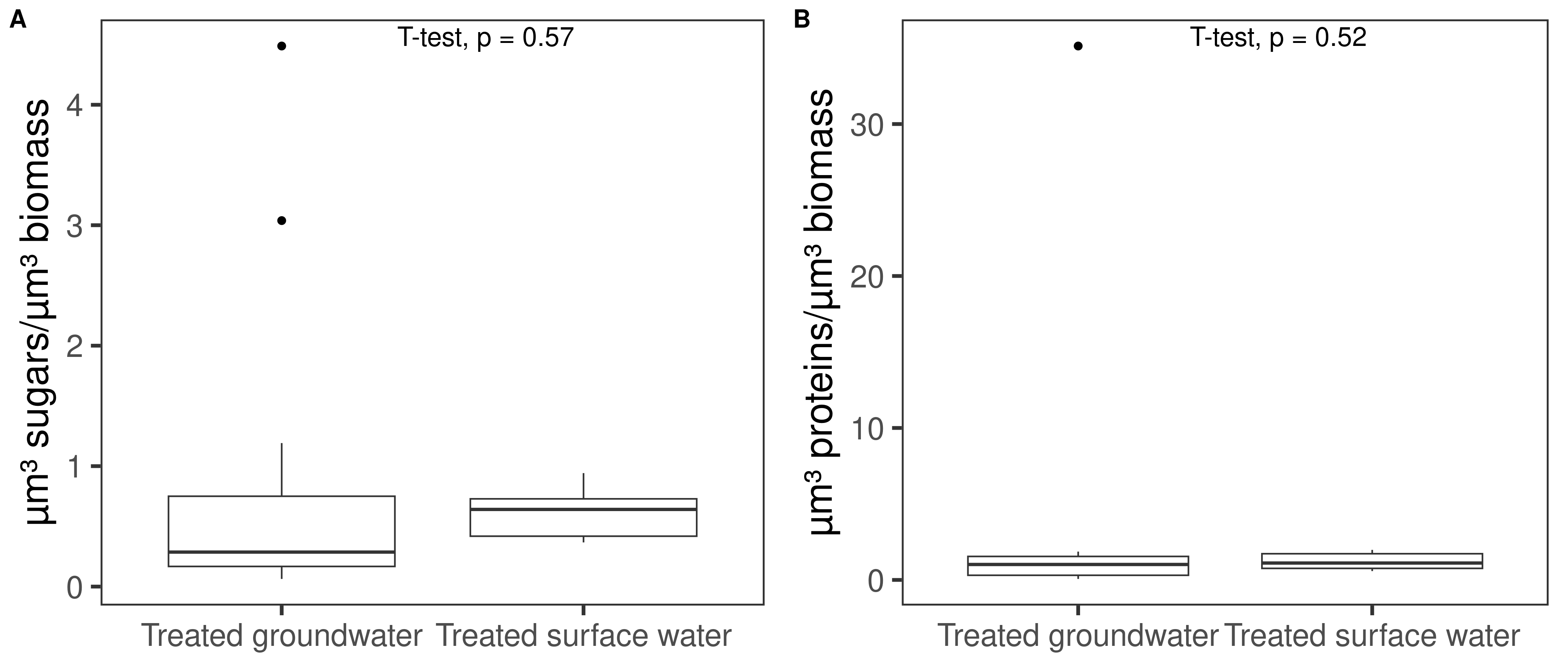

Figure A3: Ratios of **(A)** sugar content and **(B)** protein content on biomass content. Sugars, proteins and DNA were stained with Sypro Ruby, Concanavalin A and DAPI, respectively. CLSM and comstat2 were used to calculate biovolumes.

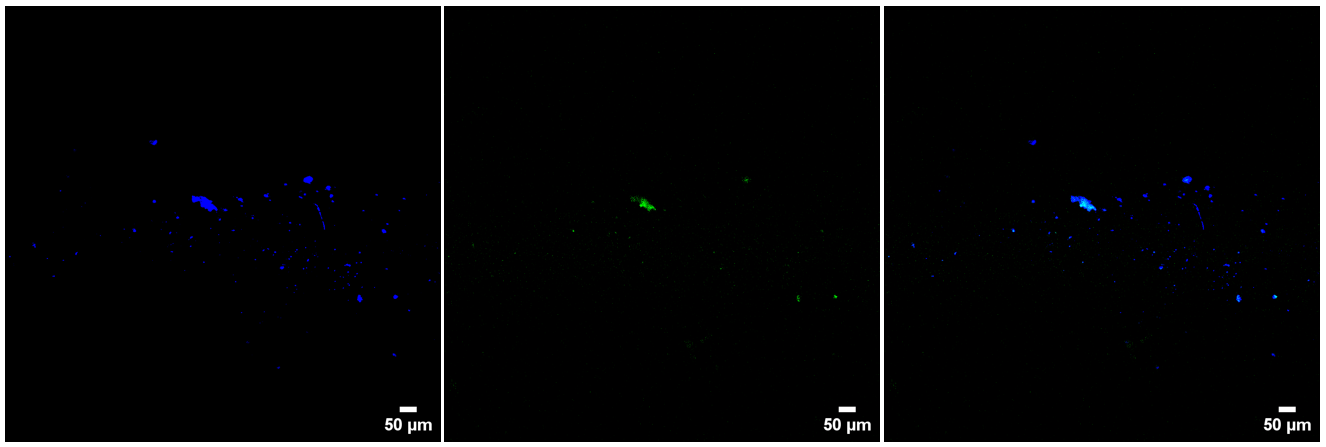

Figure A4: Confocal microscopy images (10x objective) of the 1st invasion experiment (8 days after the spike) with Serratia fonticola for the treated surface water biofilm. From left to right: biofilms stained with DAPI, the GFP-labelled coliforms, an overlay image.

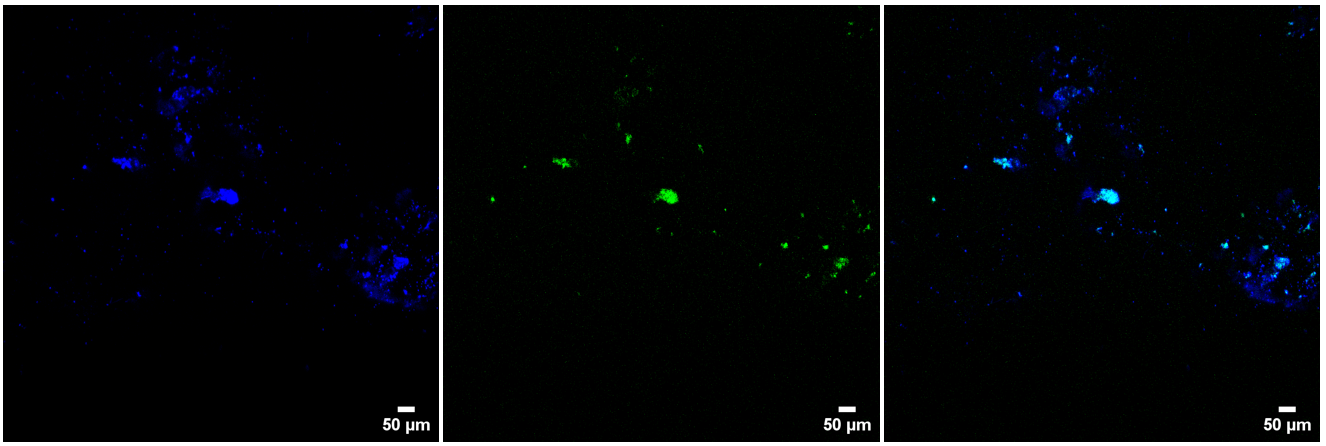

Figure A5: Confocal microscopy images (10x objective) of the 1st invasion experiment (8 days after the spike) with Serratia fonticola for the treated surface water biofilm. From left to right: biofilms stained with DAPI, the GFP-labelled coliforms, an overlay image.

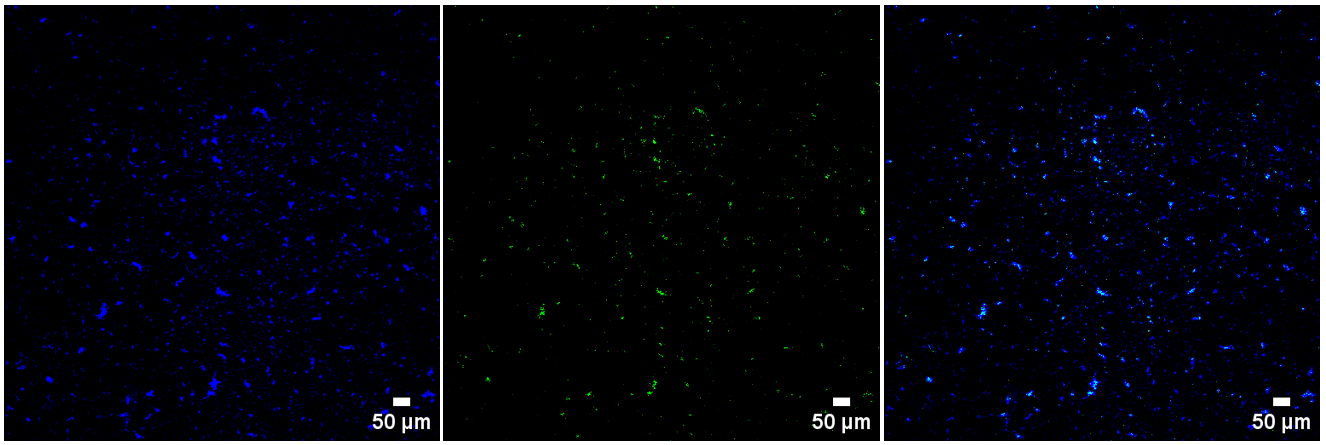

Figure A6: Confocal microscopy images (10x objective) of the 2^nd^ invasion experiment (7 days after the spike) with Serratia fonticola for the treated surface water biofilm. From left to right: biofilms stained with DAPI, the GFP-labelled coliforms, an overlay image.

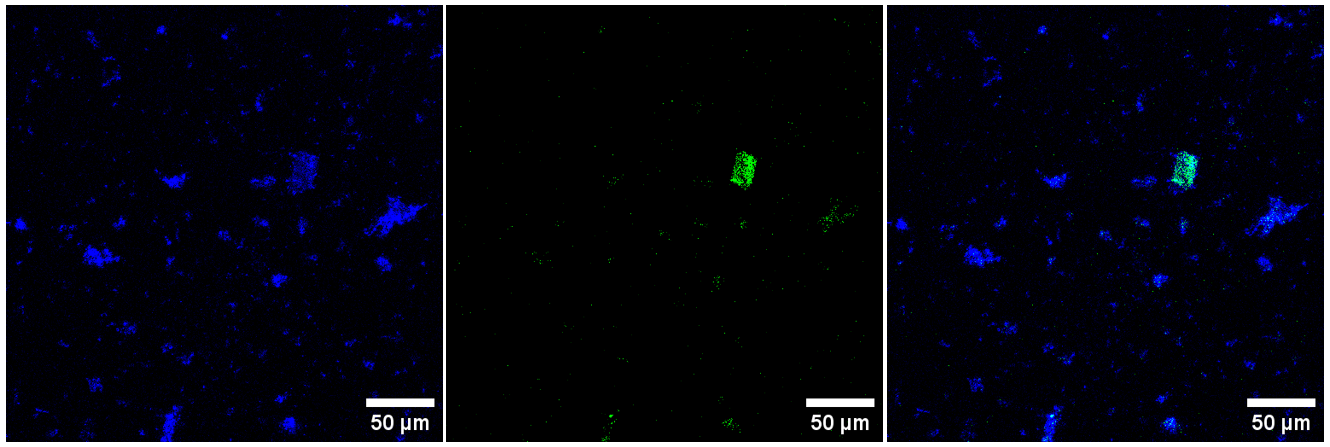

Figure A7: Confocal microscopy images (40x objective) of the 2^nd^ invasion experiment (7 days after the spike) with Serratia fonticola for the treated surface water biofilm. From left to right: biofilms stained with DAPI, the GFP-labelled coliforms, an overlay image.

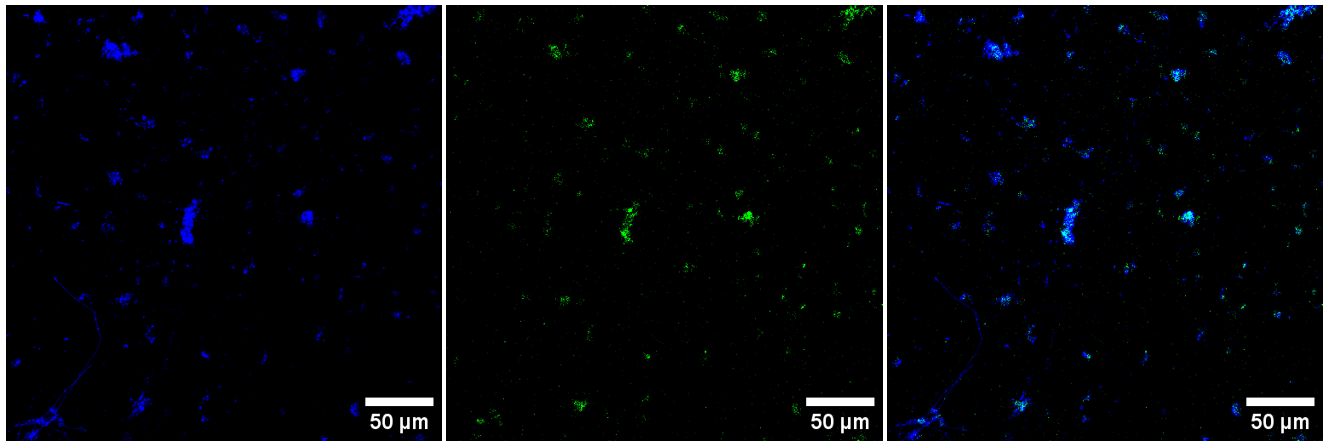

Figure A8: Confocal microscopy images (40x objective) of the 2^nd^ invasion experiment (7 days after the spike) with Serratia fonticola for the treated surface water biofilm. From left to right: biofilms stained with DAPI, the GFP-labelled coliforms, an overlay image.

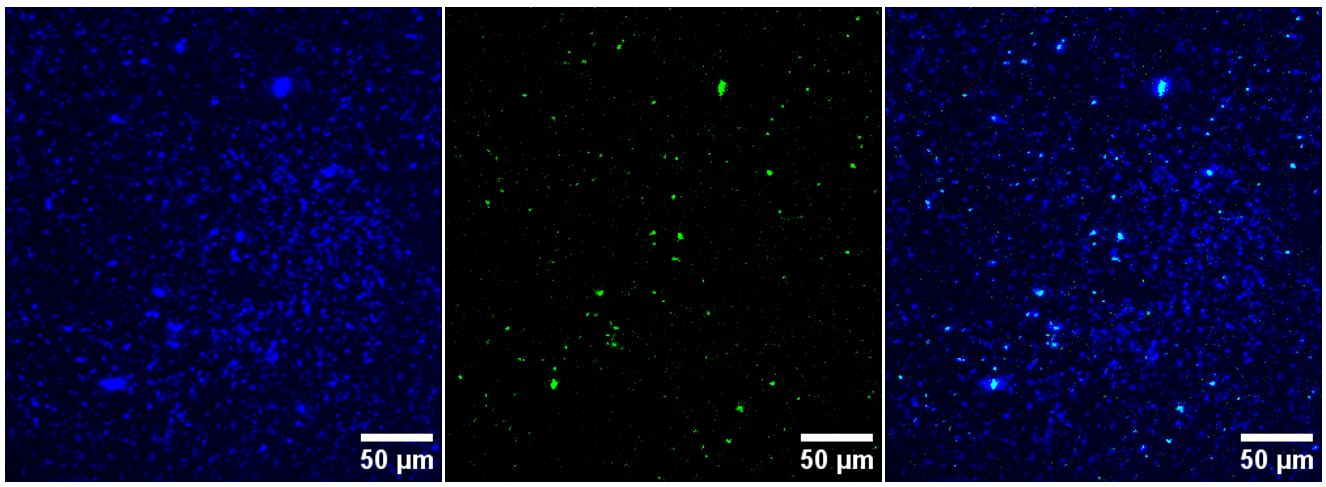

Figure A9: Confocal microscopy images (10x objective) of the 1^st^ invasion experiment (8 days after the spike) with Serratia fonticola for the treated groundwater biofilm. From left to right: biofilms stained with DAPI, the GFP-labelled coliforms, an overlay image.

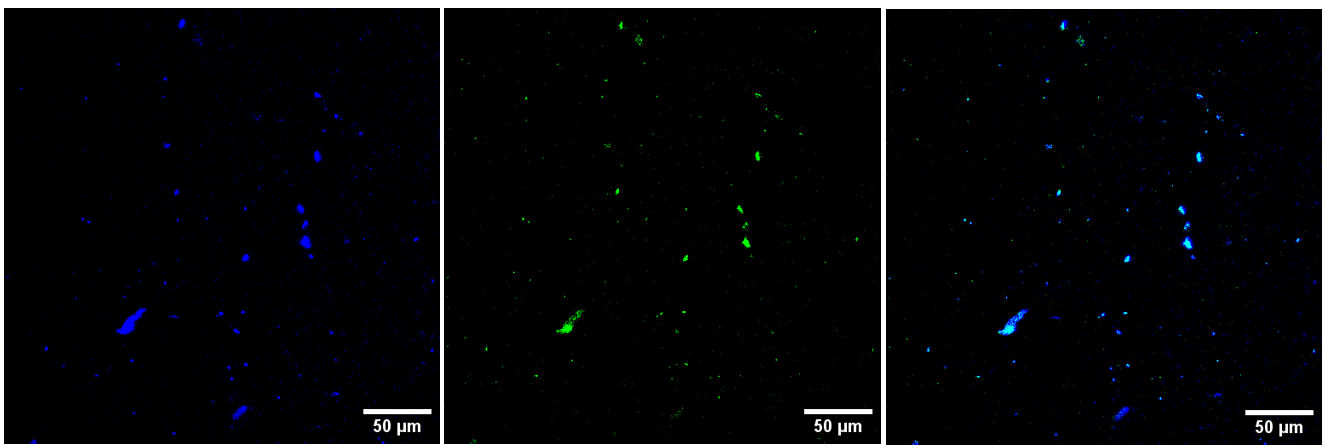

Figure A10: Confocal microscopy images (40x objective) of the 1^st^ invasion experiment (8 days after the spike) with Serratia fonticola for the treated groundwater biofilm. From left to right: biofilms stained with DAPI, the GFP-labelled coliforms, an overlay image.

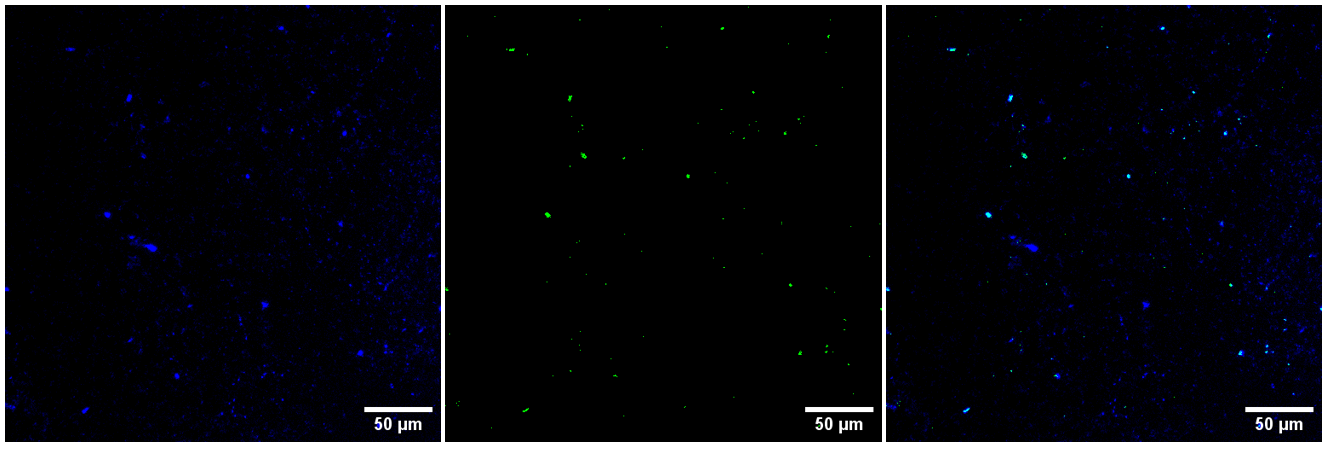

Figure A11: Confocal microscopy images (40x objective) of the 1^st^ invasion experiment (8 days after the spike) with Serratia fonticola for the treated groundwater biofilm. From left to right: biofilms stained with DAPI, the GFP-labelled coliforms, an overlay image.

*Table A1: Information of both sampling locations and the average ± standard deviation of the water quality characteristics over the duration of the experiment (17 months). Water quality parameters*^[1]^*were measured by the respective drinking water providers and averages and standard deviations are calculated using Table A5 and A6. The total organic carbon (TOC) concentration was measured at the end of the experiment*^[2]^*.*

|  | **Monitor 1** | **Monitor 2** |
| --- | --- | --- |
| Place | Water reservoir | Water tower |
| Water source | Groundwater | Surface water |
| Treatment steps | Aeration, pH correction (Ca(OH)_2_), decantation, sand filtration, storage | Aeration, pH correction (NaOH), decantation,  sand filtration, storage, disinfection (NaOCl + UV) |
| Aluminium (µg/L) | 0.18 ± 0.56 | 42.62 ± 7.77 |
| Ammonium (mg NH4/L) | 0.00 ± 0.00 | 0.02 ± 0.01 |
| Calcium (mg/L) | 47.64 ± 2.35 | 61.09 ± 10.53 |
| Total coliforms (/100 mL) | 0.16 ± 0.82 | 0.00 ± 0.00 |
| Enterococci (/100 mL) | 0.00 ± 0.00 | 0.00 ± 0.00 |
| Escherichia coli (/100 mL) | 0.00 ± 0.00 | 0.00 ± 0.00 |
| Conductivity (µS/cm) | 318.11 ± 14.83 | 459.32 ± 61.37 |
| Iron (mg/L) | 0.00 ± 0.00 | 0.00 ± 0.00 |
| Lead (µg/L) | 0.00 ± 0.00 | 0.12 ± 0.18 |
| Magnesium (mg/L) | 8.36 ± 0.52 | 7.47 ± 1.46 |
| Manganese (µg/L) | 1.13 ± 0.99 | 0.55 ± 1.03 |
| Nitrate (mg NO3/L) | 1.33 ± 0.16 | 9.94 ± 2.54 |
| Nitrite (mg NO2/L) | 0.00 ± 0.00 | 0.00 ± 0.00 |
| pH | 7.70 ± 0.12 | 7.95 ± 0.08 |
| Total colonies 22°C (/mL) | 7.66 ± 28 | 2.82 ± 5.52 |
| Temperature (°C) | 12.08 ± 0.78 | 12.30 ± 5.01 |
| Hardness (°F) | 15.32 ± 0.75 | 18.38 ± 3.16 |
| Free Cl residuals (µg/L) | 0.00 ± 0.00 | 7.65 ± 9.72 |
| TOC (ppb) | 2972.5 ± 4.33 | 2080 ± 7.07 |

**Table A2:** Experimental design detailing timepoints, specific experiments, number of rings analyzed, and types of analyses conducted. To perform FCM, ATP, 16S sequencing, and qPCR, a ring was placed in 10 mL of autoclaved tap water and subjected to sonication (3 cycles of 2 minutes each). Total cell counts (TCC) of the biofilms are reported as an average ± standard deviation before each experiment.

| **Timepoint (months)** | **Experiment** | **Rings taken for analysis** | **Type of analysis** | **Biofilm treated groundwater**  **TCC (cells/cm²)** | **Biofilm treated surface water**  **TCC (cells/cm²)** |
| --- | --- | --- | --- | --- | --- |
| 1 | Growing biofilm | / | / | / | / |
| 11 | Growing biofilm | 4 | FCM, ATP, 16S | (1.01 ± 0.04) x 10^7^ | (5.96 ± 1.01) x 10^5^ |
|  |  | 3 | CLSM (DAPI) | / | / |
| 11 | Growing biofilm | 6 | CLSM: test EPS staining | / | / |
| 17 | Growing biofilm | 4 | FCM, ATP, 16S | (7.46 ± 0.58) x 10^6^ | (5.23 ± 3.68) x 10^5^ |
|  |  | 3 | CLSM (DAPI, EPS) | / | / |
| 18 | Changing conditions: LSI = -0.50 | Hour 0: 2 | FCM | (2.70 ± 0.80) x 10^7^ | (1.25 ± 0.34) x 10^6^ |
| 18 | Changing conditions: LSI = 0.30 | Hour 0: 2 | FCM | (1.23 ± 0.12) x 10^7^ | (2.36 ± 0.11) x 10^6^ |
| 19 | Changing conditions: HOCl addition | Hour 0: 2 | FCM | (1.76 ± 0.44) x 10^7^ | (2.92 ± 0.08) x 10^6^ |
| 20 | Invasion (1^st^ experiment) | Day 0: 2 | FCM, qPCR,  CLSM (DAPI) | (1.57 ± 0.06) x 10^7^ | (1.05 ± 0.01) x 10^6^ |
|  |  | Day 4: 2 | qPCR, CLSM (DAPI) | / | / |
|  |  | Day 7: 1 | CLSM (DAPI) | / | / |
| 24 | Invasion (2^nd^ experiment) | Day 0: 2 | FCM, qPCR  CLSM (DAPI) | (7.56 ± 0.09) x 10^6^ | (6.87 ± 0.02) x 10^5^ |
|  |  | Day 8: 1 | qPCR, CLSM (DAPI) | / | / |

Table A3: Characterization of the biofilms using different techniques after 11 and 17 months. Results are presented as an average ± standard deviation. Statistics are done between the different timepoints using the Wilcoxon rank sum test and the t test indicated with a ‘t’. Bray-curtis dissimilarity and ANOSIM were used to calculate statistics for the 16S rRNA sequencing data. Biomass and community composition were not significantly different over time for both biofilms showing that the biofilms were mature over time.

| **Technique** | **Parameter** | **After 11 months** | **After 17 months** | **P value** | **After 11 months** | **After 17 months** | **P value** |
| --- | --- | --- | --- | --- | --- | --- | --- |
| **Water type** | **Water type** | **Groundwater (not disinfected)** | **Groundwater (not disinfected)** |  | **Surface water (chlorinated)** | **Surface water (chlorinated)** |  |
| **Flow cytometry** | **Total cell counts (cells/cm²)** | (1.01 ± 0.04) x 10^7^ | (7.46 ± 0.58) x 10^6^ | 0.01061^t^ | (5.96 ± 1.01) x 10^5^ | (5.23 ± 3.68) x 10^5^ | 0.8121 |
| **ATP analysis** | **ATP content (pg/cm²)** | 411.34 ± 52.58 | 896.54 ± 72.11 | 3.275e-10^t^ | 51.86 ± 3.22 | 77.26 ± 41.48 | 0.1104^t^ |
| **CLSM** | **Roughness (Ra)** | 1.25 ± 0.39 | 0.78 ± 0.14 | 0.0295^t^ | 1.97 ± 0.03 | 1.68 ± 0.04 | 0.004615^t^ |
| **CLSM** | **Average thickness (µm)** | 15.78 ± 5.89 | 4.95 ± 2.62 | 0.0008926^t^ | 0.94 ± 0.57 | 1.31 ± 0.33 | 0.3569^t^ |
| **CLSM** | **Biomass (µm³/µm²)** | 5.65 ± 2.23 | 4.10 ± 3.07 | 0.511^t^ | 0.57 ± 0.45 | 0.25 ± 0.10 | 0.2906^t^ |
| **CLSM** | **Maximum thickness (µm)** | 86.80 ± 27.72 | 41. 95 ± 37.05 | 0.06943 | 127.6 ± 57.71 | 19.5 ± 3.04 | 0.009858 |
| **NGS** | **Composition (-)** | / | / | 0.333 | / | / | 0.6667 |

Table A4: Biofilm detachment was evaluated by changing the operational water quality characteristics. The applied conditions were compared with a blank conditions and a linear regression model was calculated. Statistics on the residuals of the regression models were done with a t test. In addition, significance of the differences between the slopes was considered. Here, p-values were calculated by comparing the observed difference of the slopes to the distribution of bootstrapped differences using resampled datasets. This provided an estimate of the probability of observing a difference as extreme as (or more extreme than) the observed difference, assuming the null hypothesis (no difference in slopes) is true.

|  | P value of the residuals of the regression model | Significance between the slopes (bootstrapping) |
| --- | --- | --- |
| Groundwater, HOCl addition | 0.1318 | 0.473 |
| Groundwater, LSI = -0.50 | 0.1418 | 0.494 |
| Groundwater, LSI = 0.30 | 0.0248 | 0.651 |
| Surface water, HOCl addition | 0.1176 | 0.532 |
| Surface water, LSI = -0.50 | 0.0312 | 0.479 |
| Surface water, LSI = 0.30 | 0.1778 | 0.764 |

Table A5: Raw water quality measurements of treated groundwater without chlorination over the duration of the experiment (17 months), measured^1^^[3]^ by the respective drinking water providers.

| **Date** | **Al (µg/L)** | **NH4+(mg NH4/L)** | **Calcium (mg/L)** | **Total coliforms (/100 mL)** | **Enterococci (/100 mL)** | **Escherichia coli (/100 mL)** | **Conductivity (µS/cm)** | **Fe(mg/L)** | **Pb (µg/L)** | **Mg (mg/L)** | **Mn (µg/L)** | **NO3- (mg NO3/L)** | **NO2- (mg NO2/L)** | **pH** | **Total colonies 22°C (/mL)** | **Temperature (°C)** | **Hardness (°F)** | **Free Cl residuals (µg/L)** |
| --- | --- | --- | --- | --- | --- | --- | --- | --- | --- | --- | --- | --- | --- | --- | --- | --- | --- | --- |
| 14/12/2020 | 0.00 | 0 | 43.78 | 0 | 0 | 0 | 302.70 | 0 | 0 | 7.66 | 0.72 | 1.15 | 0 | 7.83 | 5 | 11.60 | 14.07 | 0 |
| 21/12/2020 | 0.00 | 0 | 43.01 | 0 | 0 | 0 | 300.00 | 0 | 0 | 7.90 | 0.82 | 1.35 | 0 | 7.68 | 0 | 12.50 | 13.98 | 0 |
| 28/12/2020 | 0.00 | 0 | 46.99 | 0 | 0 | 0 | 316.22 | 0 | 0 | 8.22 | 4.74 | 1.42 | 0 | 7.65 | 16 | 11.70 | 15.10 | 0 |
| 4/1/2021 | 0.00 | 0 | 45.63 | 0 | 0 | 0 | 310.81 | 0 | 0 | 8.08 | 0.94 | 1.26 | 0 | 7.82 | 3 | 11.60 | 14.70 | 0 |
| 11/1/2021 | 0.00 | 0 | 49.19 | 0 | 0 | 0 | 327.93 | 0 | 0 | 9.06 | 6.87 | 1.49 | 0 | 7.76 | 52 | 11.10 | 15.99 | 0 |
| 18/1/2021 | 2.11 | 0 | 49.91 | 0 | 0 | 0 | 327.93 | 0 | 0 | 9.23 | 2.13 | 1.53 | 0 | 7.65 | 0 | 11.40 | 16.25 | 0 |
| 25/1/2021 | 0.00 | 0 | 46.78 | 0 | 0 | 0 | 306.31 | 0 | 0 | 8.37 | 1.63 | 1.18 | 0 | 7.74 | 9 | 11.20 | 15.11 | 0 |
| 1/2/2021 | 0.00 | 0 | 46.77 | 0 | 0 | 0 | 301.80 | 0 | 0 | 8.45 | 1.79 | 1.29 | 0 | 7.67 | 0 | 11.70 | 15.14 | 0 |
| 8/2/2021 | 0.00 | 0 | 49.03 | 0 | 0 | 0 | 312.61 | 0 | 0 | 8.62 | 1.48 | 1.20 | 0 | 7.69 | 0 | 8.30 | 15.77 | 0 |
| 15/2/2021 | 0.00 | 0 | 45.46 | 0 | 0 | 0 | 314.41 | 0 | 0 | 8.23 | 1.14 | 1.26 | 0 | 7.75 | 3 | 11.80 | 14.73 | 0 |
| 22/2/2021 | 0.00 | 0 | 49.55 | 0 | 0 | 0 | 336.04 | 0 | 0 | 8.57 | 1.40 | 1.31 | 0 | 7.69 | 0 | 11.80 | 15.88 | 0 |
| 1/3/2021 | 0.00 | 0 | 48.30 | 0 | 0 | 0 | 323.42 | 0 | 0 | 8.42 | 1.25 | 1.31 | 0 | 7.66 | 2 | 11.60 | 15.51 | 0 |
| 8/3/2021 | 0.00 | 0 | 46.67 | 0 | 0 | 0 | 320.72 | 0 | 0 | 7.89 | 0.79 | 1.35 | 0 | 7.77 | 1 | 11.60 | 14.88 | 0 |
| 15/3/2021 | 0.00 | 0 | 44.16 | 0 | 0 | 0 | 300.00 | 0 | 0 | 7.68 | 0.86 | 1.25 | 0 | 7.76 | 2 | 11.60 | 14.18 | 0 |
| 22/3/2021 | 0.00 | 0 | 48.95 | 0 | 0 | 0 | 326.13 | 0 | 0 | 8.77 | 0.91 | 1.34 | 0 | 7.71 | 6 | 11.70 | 15.81 | 0 |
| 29/3/2021 | 0.00 | 0 | 46.22 | 0 | 0 | 0 | 306.31 | 0 | 0 | 8.20 | 0.83 | 1.16 | 0 | 7.71 | 33 | 11.90 | 14.90 | 0 |
| 6/4/2021 | 0.00 | 0 | 46.57 | 0 | 0 | 0 | 296.40 | 0 | 0 | 8.36 | 0.64 | 1.33 | 0 | 7.72 | 47 | 11.80 | 15.05 | 0 |
| 12/4/2021 | 0.00 | 0 | 47.32 | 0 | 0 | 0 | 310.81 | 0 | 0 | 8.09 | 0.77 | 1.18 | 0 | 7.69 | 1 | 11.90 | 15.13 | 0 |
| 19/4/2021 | 0.00 | 0 | 47.15 | 0 | 0 | 0 | 336.94 | 0 | 0 | 8.26 | 0.81 | 1.44 | 0 | 7.80 | 3 | 11.70 | 15.16 | 0 |
| 26/4/2021 | 0.00 | 0 | 43.35 | 0 | 0 | 0 | 302.70 | 0 | 0 | 7.48 | 0.61 | 1.17 | 0 | 7.86 | 2 | 11.90 | 13.89 | 0 |
| 3/5/2021 | 0.00 | 0 | 49.31 | 0 | 0 | 0 | 326.13 | 0 | 0 | 8.38 | 0.70 | 1.35 | 0 | 7.85 | 7 | 12.40 | 15.75 | 0 |
| 10/5/2021 | 0.00 | 0 | 45.96 | 0 | 0 | 0 | 318.92 | 0 | 0 | 8.51 | 1.08 | 1.38 | 0 | 7.61 | 1 | 12.40 | 14.96 | 0 |
| 17/5/2021 | 0.00 | 0 | 48.19 | 0 | 0 | 0 | 314.41 | 0 | 0 | 9.28 | 1.23 | 1.51 | 0 | 7.57 | 23 | 12.40 | 15.84 | 0 |
| 25/5/2021 | 0.00 | 0 | 48.36 | 0 | 0 | 0 | 340.54 | 0 | 0 | 8.75 | 0.92 | 1.39 | 0 | 7.70 | 1 | 12.30 | 15.66 | 0 |
| 31/5/2021 | 0.00 | 0 | 49.78 | 0 | 0 | 0 | 334.23 | 0 | 0 | 8.88 | 0.83 | 1.58 | 0 | 7.71 | 0 | 12.60 | 16.07 | 0 |
| 7/6/2021 | 0.00 | 0 | 45.01 | 0 | 0 | 0 | 298.20 | 0 | 0 | 6.48 | 0.00 | 1.07 | 0 | 8.09 | 1 | 12.60 | 13.89 | 0 |
| 14/6/2021 | 0.00 | 0 | 55.44 | 0 | 0 | 0 | 358.56 | 0 | 0 | 8.82 | 1.04 | 1.61 | 0 | 7.72 | 7 | 13.10 | 17.46 | 0 |
| 21/6/2021 | 0.00 | 0 | 47.65 | 0 | 0 | 0 | 305.41 | 0 | 0 | 7.88 | 2.87 | 1.21 | 0 | 7.70 | 223 | 13.00 | 15.12 | 0 |
| 28/6/2021 | 0.00 | 0 | 48.62 | 0 | 0 | 0 | 319.82 | 0 | 0 | 8.71 | 0.00 | 1.43 | 0 | 7.64 | 0 | 13.00 | 15.71 | 0 |
| 5/7/2021 | 0.00 | 0 | 46.12 | 0 | 0 | 0 | 290.09 | 0 | 0 | 8.35 | 1.03 | 1.19 | 0 | 7.71 | 2 | 12.90 | 14.94 | 0 |
| 12/7/2021 | 0.00 | 0 | 48.66 | 0 | 0 | 0 | 329.73 | 0 | 0 | 8.61 | 0.67 | 1.40 | 0 | 7.69 | 2 | 13.20 | 15.68 | 0 |
| 19/7/2021 | 0.00 | 0 | 47.19 | 0 | 0 | 0 | 319.82 | 0 | 0 | 8.45 | 0.00 | 1.18 | 0 | 7.76 | 0 | 13.10 | 15.24 | 0 |
| 26/7/2021 | 0.00 | 0 | 46.26 | 0 | 0 | 0 | 319.82 | 0 | 0 | 8.22 | 0.74 | 1.22 | 0 | 7.63 | 3 | 13.10 | 14.92 | 0 |
| 2/8/2021 | 0.00 | 0 | 48.03 | 0 | 0 | 0 | 309.01 | 0 | 0 | 8.54 | 0.52 | 1.47 | 0 | 7.61 | 0 | 13.00 | 15.49 | 0 |
| 9/8/2021 | 3.97 | 0 | 46.09 | 0 | 0 | 0 | 304.50 | 0 | 0 | 7.87 | 0.00 | 1.34 | 0 | 7.56 | 1 | 12.60 | 14.73 | 0 |
| 16/8/2021 | 0.00 | 0 | 51.76 | 0 | 0 | 0 | 341.44 | 0 | 0 | 8.65 | 0.59 | 1.39 | 0 | 7.68 | 0 | 13.00 | 16.47 | 0 |
| 23/8/2021 | 0.00 | 0 | 46.14 | 0 | 0 | 0 | 309.01 | 0 | 0 | 8.32 | 0.56 | 1.39 | 0 | 7.70 | 0 | 13.10 | 14.93 | 0 |
| 30/8/2021 | 0.00 | 0 | 48.11 | 0 | 0 | 0 | 315.32 | 0 | 0 | 8.74 | 0.62 | 1.54 | 0 | 7.66 | 2 | 12.90 | 15.59 | 0 |
| 6/9/2021 | 0.00 | 0 | 48.72 | 0 | 0 | 0 | 327.03 | 0 | 0 | 8.78 | 0.78 | 1.52 | 0 | 7.65 | 2 | 13.20 | 15.76 | 0 |
| 13/9/2021 | 0.00 | 0 | 44.66 | 0 | 0 | 0 | 310.81 | 0 | 0 | 8.10 | 0.77 | 1.15 | 0 | 7.61 | 1 | 13.20 | 14.47 | 0 |
| 20/9/2021 | 0.00 | 0 | 51.66 | 0 | 0 | 0 | 348.65 | 0 | 0 | 8.65 | 0.51 | 1.51 | 0 | 7.65 | 0 | 13.00 | 16.44 | 0 |
| 27/9/2021 | 0.00 | 0 | 46.77 | 0 | 0 | 0 | 300.90 | 0 | 0 | 8.44 | 0.00 | 1.15 | 0 | 7.69 | 1 | 13.00 | 15.14 | 0 |
| 4/10/2021 | 0.00 | 0 | 49.39 | 0 | 0 | 0 | 330.63 | 0 | 0 | 9.21 | 0.56 | 1.35 | 0 | 7.61 | 0 | 12.90 | 16.10 | 0 |
| 11/10/2021 | 0.00 | 0 | 49.95 | 0 | 0 | 0 | 343.24 | 0 | 0 | 8.63 | 0.57 | 1.41 | 0 | 7.71 | 1 | 12.50 | 16.01 | 0 |
| 18/10/2021 | 0.00 | 0 | 49.22 | 0 | 0 | 0 | 345.05 | 0 | 0 | 8.04 | 0.59 | 1.38 | 0 | 7.76 | 1 | 12.30 | 15.58 | 0 |
| 25/10/2021 | 0.00 | 0 | 45.39 | 0 | 0 | 0 | 301.80 | 0 | 0 | 8.09 | 0.78 | 1.20 | 0 | 7.73 | 2 | 12.20 | 14.65 | 0 |
| 4/11/2021 | 0.00 | 0 | 51.92 | 0 | 0 | 0 | 324.32 | 0 | 0 | 8.14 | 0.82 | 1.31 | 0 | 7.73 | 0 | 12.40 | 16.30 | 0 |
| 10/11/2021 | 0.00 | 0 | 48.49 | 3 | 0 | 0 | 319.82 | 0 | 0 | 8.75 | 0.77 | 1.34 | 0 | 7.77 | 0 | 12.30 | 15.69 | 0 |
| 29/11/2021 | 0.00 | 0 | 47.04 | 6 | 0 | 0 | 326.13 | 0 | 0 | 7.95 | 0.57 |  | 0 | 7.82 | 0 | 12.10 | 15.00 | 0 |
| 6/12/2021 | 0.00 | 0 | 49.77 | 0 | 0 | 0 | 318.02 | 0 | 0 | 8.77 | 0.75 | 1.50 | 0 | 7.77 | 1 | 11.90 | 16.02 | 0 |
| 13/12/2021 | 0.00 | 0 | 49.79 | 0 | 0 | 0 | 320.72 | 0 | 0 | 8.81 | 1.49 | 1.38 | 0 | 7.83 | 0 | 12.00 | 16.04 | 0 |
| 20/12/2021 | 0.00 | 0 | 47.49 | 0 | 0 | 0 | 322.52 | 0 | 0 | 8.60 | 1.71 | 1.43 | 0 | 7.57 | 2 | 10.30 | 15.38 | 0 |
| 27/12/2021 | 0.00 | 0 | 45.18 | 0 | 0 | 0 | 338.74 | 0 | 0 | 7.89 | 1.78 | 1.06 | 0 | 7.54 | 3 | 11.70 | 14.51 | 0 |
| 1/3/2022 | 0.00 | 0 | 47.60 | 0 | 0 | 0 | 300.90 | 0 | 0 | 8.10 | 0.78 | 1.38 | 0 | 7.80 | 0 | 11.60 | 15.20 | 0 |
| 1/10/2022 | 0.00 | 0 | 44.79 | 0 | 0 | 0 | 327.93 | 0 | 0 | 8.46 | 1.43 | 1.24 | 0 | 7.80 | 0 | 12.00 | 14.65 | 0 |
| 1/17/2022 | 0.00 | 0 | 44.40 | 0 | 0 | 0 | 313.51 | 0 | 0 | 7.23 | 1.43 | 1.09 | 0 | 7.69 | 1 | 11.60 | 14.05 | 0 |
| 1/31/2022 | 0.58 | 0 | 46.27 | 0 | 0 | 0 | 308.11 | 0 | 0 | 8.55 | 1.71 | 1.27 | 0 | 7.83 | 4 | 11.70 | 15.06 | 0 |
| 2/7/2022 | 0.00 | 0 | 46.82 | 0 | 0 | 0 | 299.10 | 0 | 0 | 8.22 | 1.06 | 1.42 | 0 | 7.60 | 2 | 11.60 | 15.06 | 0 |
| 2/14/2022 | 0.49 | 0 | 44.70 | 0 | 0 | 0 | 333.33 | 0 | 0 | 7.89 | 0.97 | 1.23 | 0 | 7.80 | 1 | 11.60 | 14.39 | 0 |
| 2/21/2022 | 0.46 | 0 | 47.92 | 0 | 0 | 0 | 300.00 | 0 | 0 | 7.69 | 1.55 | 1.22 | 0 | 7.78 | 3 | 11.60 | 15.11 | 0 |
| 2/28/2022 | 0.31 | 0 | 47.79 | 1 | 0 | 0 | 328.83 | 0 | 0 | 8.20 | 2.02 | 1.41 | 0 | 7.71 | 2 | 11.80 | 15.29 | 0 |
| 3/7/2022 | 0.40 | 0 | 44.34 | 0 | 0 | 0 | 334.23 | 0 | 0 | 7.85 | 1.64 | 1.33 | 0 | 7.78 | 2 | 11.50 | 14.29 | 0 |
| 3/14/2022 | 0.40 | 0 | 47.81 | 0 | 0 | 0 | 299.10 | 0 | 0 | 8.22 | 1.67 | 1.40 | 0 | 7.66 | 18 | 11.50 | 15.30 | 0 |
| 3/21/2022 | 0.75 | 0 | 50.80 | 0 | 0 | 0 | 296.40 | 0 | 0 | 8.88 | 0.83 | 1.39 | 0 | 7.73 | 5 | 11.60 | 16.32 | 0 |
| 3/28/2022 | 0.35 | 0 | 51.38 | 0 | 0 | 0 | 332.43 | 0 | 0 | 8.79 | 0.89 | 1.51 | 0 | 7.68 | 9 | 11.70 | 16.43 | 0 |
| 4/4/2022 | 0.50 | 0 | 52.98 | 0 | 0 | 0 | 318.02 | 0 | 0 | 10.14 | 1.60 | 1.62 | 0 | 7.68 | 1 | 11.90 | 17.38 | 0 |
| 4/11/2022 | 0.56 | 0 | 47.61 | 0 | 0 | 0 | 323.42 | 0 | 0 | 8.13 | 1.02 | 1.29 | 0 | 7.55 | 1 | 11.50 | 15.22 | 0 |
| 4/19/2022 | 1.71 | 0 | 47.56 | 1 | 0 | 0 | 302.70 | 0 | 0 | 8.01 | 0.96 | 1.11 | 0 | 7.02 | 0 | 12.00 | 15.16 | 0 |

Table A6: Raw water quality measurements of treated surface water with chlorination over the duration of the experiment (17 months), measured^1^^[4]^by the respective drinking water providers.

| **Date** | **Al (µg/L)** | **NH4+(mg NH4/L)** | **Calcium (mg/L)** | **Total coliforms (/100 mL)** | **Enterococci (/100 mL)** | **Escherichia coli (/100 mL)** | **Conductivity (µS/cm)** | **Fe(mg/L)** | **Pb (µg/L)** | **Mg (mg/L)** | **Mn (µg/L)** | **NO3- (mg NO3/L)** | **NO2- (mg NO2/L)** | **pH** | **Total colonies 22°C (/mL)** | **Temperature (°C)** | **Hardness (°F)** | **Free Cl residuals (µg/L)** |
| --- | --- | --- | --- | --- | --- | --- | --- | --- | --- | --- | --- | --- | --- | --- | --- | --- | --- | --- |
| 17/11/2020 | 36.01 | 0.02 | 70.73 | 0 | 0 | 0 | 638 | 1.22 | 0.08 | 9.91 | 0.10 | 6.15 | 0.00 | 7.87 | 0 | 12.90 | 21.81 | 20 |
| 1/12/2020 | 44.86 | 0.01 | 76.16 | 0 | 0 | 0 | 652 | 1.38 | 0.10 | 10.42 | 0.10 | 7.48 | 0.00 | 7.90 | 0 | 11.30 | 23.38 | 0 |
| 15/12/2020 | 32.54 | 0.02 | 74.07 | 0 | 0 | 0 | 395 | 1.33 | 0.05 | 9.29 | 0.17 | 9.15 | 0.00 | 7.86 | 4 | 9.00 | 22.39 | 0 |
| 29/12/2020 | 35.77 | 0.01 | 72.48 | 0 | 0 | 0 | 547 | 2.09 | 0.05 | 8.56 | 0.17 | 10.47 | 0.00 | 7.89 | 0 | 8.80 | 21.69 | 0 |
| 12/1/2021 | 30.38 | 0.02 | 70.08 | 0 | 0 | 0 | 506 | 2.52 | 0.05 | 7.78 | 0.20 | 11.16 | 0.00 | 7.92 | 0 | 7.10 | 20.76 | 10 |
| 26/1/2021 |  | 0.02 |  | 0 | 0 | 0 | 488 |  |  |  |  | 11.41 | 0.00 | 7.93 | 0 | 6.40 |  | 40 |
| 9/2/2021 | 37.03 | 0.02 | 62.32 | 0 | 0 | 0 | 464 | 2.30 | 0.02 | 6.94 | 0.16 | 11.51 | 0.00 | 7.93 | 0 | 6.60 | 18.47 | 20 |
| 23/2/2021 | 36.82 | 0.04 | 54.01 | 0 | 0 | 0 | 420 | 2.26 | 0.03 | 6.05 | 0.17 | 12.05 | 0.01 | 8.00 | 0 | 5.60 | 16.02 | 10 |
| 9/3/2021 | 32.20 | 0.02 | 55.75 | 0 | 0 | 0 | 416 | 2.26 | 0.03 | 5.74 | 0.25 | 12.73 | 0.00 | 8.02 | 5 | 7.50 | 16.33 | 0 |
| 23/3/2021 |  | 0.03 |  | 0 | 0 | 0 | 402 |  |  |  |  | 13.21 | 0.00 | 8.05 | 0 | 8.10 |  | 20 |
| 6/4/2021 | 43.51 | 0.02 | 56.60 | 0 | 0 | 0 | 401 | 1.22 | 0.07 | 5.63 | 0.13 | 13.03 | 0.00 | 8.08 | 1 | 10.50 | 16.49 | 0 |
| 20/4/2021 | 36.43 | 0.02 | 63.40 | 0 | 0 | 0 | 433 | 0.88 | 0.05 | 5.87 | 0.16 | 13.01 | 0.00 | 8.03 | 1 | 10.10 | 18.29 | 20 |
| 3/5/2021 | 43.76 | 0.02 | 59.22 | 0 | 0 | 0 | 432 | 1.97 | 0.07 | 6.49 | 0.55 | 12.32 | 0.00 | 8.04 | 0 | 12.10 | 17.51 | 20 |
| 18/5/2021 | 50.35 | 0.02 | 59.27 | 0 | 0 | 0 | 432 | 1.44 | 0.05 | 6.63 | 0.38 | 11.48 | 0.00 | 7.95 | 0 | 13.60 | 17.58 | 10 |
| 1/6/2021 |  | 0.02 |  | 0 | 0 | 0 | 435 |  |  |  |  | 10.43 | 0.00 | 7.94 | 1 | 14.60 |  | 0 |
| 15/6/2021 | 51.54 | 0.02 | 51.66 | 0 | 0 | 0 | 430 | 1.89 | 0.12 | 6.77 | 5.60 | 8.59 | 0.00 | 7.95 | 5 | 19.20 | 15.73 | 0 |
| 29/6/2021 | 48.02 | 0.02 | 52.48 | 0 | 0 | 0 | 441 | 2.44 | 0.11 | 7.29 | 0.46 | 7.50 | 0.00 | 7.98 | 1 |  | 16.16 | 0 |
| 13/7/2021 | 52.12 | 0.02 | 52.97 | 0 | 0 | 0 | 449 | 3.80 | 0.13 | 7.51 | 0.77 | 6.74 | 0.00 | 7.97 | 4 | 20.10 | 16.37 | 0 |
| 27/7/2021 | 50.07 | 0.02 | 42.51 | 0 | 0 | 0 | 406 | 7.70 | 0.22 | 6.33 | 1.30 | 6.45 | 0.00 | 8.02 | 0 | 21.60 | 13.26 | 10 |
| 10/8/2021 | 62.51 | 0.02 | 48.35 | 0 | 0 | 0 | 411 | 3.78 | 1.01 | 6.71 | 0.54 | 5.90 | 0.00 | 8.07 | 4 | 20.50 | 14.88 | 10 |
| 24/8/2021 | 45.66 | 0.02 | 48.79 | 0 | 0 | 0 | 414 | 9.15 | 0.29 | 7.27 | 1.86 | 5.66 | 0.00 | 8.00 | 0 | 20.20 | 15.22 | 20 |
| 7/9/2021 | 40.87 | 0.02 | 45.57 | 0 | 0 | 0 | 407 | 3.21 | 0.06 | 5.91 | 0.53 | 6.04 | 0.00 | 7.98 | 2 | 20.20 | 13.86 | 0 |
| 21/9/2021 |  | 0.02 |  | 0 | 0 | 0 | 425 |  |  |  |  | 6.25 | 0.01 | 8.06 | 20 | 19.90 |  | 0 |
| 5/10/2021 | 35.16 | 0.03 | 58.31 | 0 | 0 | 0 | 449 | 7.42 | 0.24 | 6.96 | 0.88 | 6.88 | 0.00 | 7.95 | 16 | 18.20 | 17.48 | 10 |
| 19/10/2021 | 33.96 | 0.02 | 61.14 | 0 | 0 | 0 | 464 | 1.53 | 0.12 | 7.41 | 0.20 | 7.85 | 0.00 | 7.94 | 23 | 15.80 | 18.37 | 10 |
| 3/11/2021 | 45.68 | 0.01 | 64.79 | 0 | 0 | 0 | 480 | 2.65 | 0.08 | 7.71 | 0.20 | 9.18 | 0.00 | 7.98 | 2 | 14.30 | 19.41 | 0 |
| 16/11/2021 | 39.40 | 0.02 | 76.69 | 0 | 0 | 0 | 535 | 1.55 | 0.11 | 9.88 | 0.23 | 11.32 | 0.01 | 7.82 | 1 | 12.70 | 23.29 | 0 |
| 30/11/2021 |  | 0.02 |  | 0 | 0 | 0 | 494 |  |  |  |  | 10.93 | 0.00 | 8.00 | 1 | 10.60 |  | 20 |
| 14/12/2021 | 39.38 | 0.00 | 60.46 | 0 | 0 | 0 | 466 | 2.01 | 0.06 | 7.37 | 0.10 | 11.57 | 0.01 | 7.90 | 1 | 8.70 | 18.19 | 0 |
| 28/12/2021 | 28.89 | 0.00 | 91.76 | 0 | 0 | 0 | 546 | 1.83 | 0.07 | 11.41 | 0.28 | 12.64 | 0.00 | 7.68 | 1 | 8.30 | 27.69 | 0 |
| 11/1/2022 | 42.35 | 0.01 | 67.29 | 0 | 0 | 0 | 466 | 2.99 | 0.05 | 8.65 | 0.15 | 12.29 | 0.00 | 7.81 | 0 | 8.70 | 20.43 | 10 |
| 25/1/2022 | 31.41 | 0.02 | 55.79 | 0 | 0 | 0 | 429 | 1.94 | 0.04 | 6.57 | 0.11 | 12.06 | 0.00 | 7.96 | 0 | 7.30 | 16.69 | 0 |
| 8/2/2022 | 33.21 | 0.02 | 59.10 | 0 | 0 | 0 | 424 | 2.00 | 0.04 | 6.85 | 0.10 | 12.03 | 0.01 | 7.94 | 0 | 7.50 | 17.63 | 0 |
| 22/2/2022 | 38.24 | 0.01 | 59.89 | 0 | 0 | 0 | 420 | 1.96 | 0.07 | 6.68 | 0.01 | 12.51 | 0.00 | 8.02 | 3 | 7.90 | 17.75 | 0 |

1. emis, “Compendium Voor de Monsterneming, Meting En Analyse van Water.”
2. Waegenaar et al., “Insects in Water Towers.”
3. ^1^ emis, “Compendium Voor de Monsterneming, Meting En Analyse van Water.”
4. ^1^ emis, “Compendium Voor de Monsterneming, Meting En Analyse van Water.”
